# Short-term facilitation of long-range corticocortical synapses revealed by selective optical stimulation

**DOI:** 10.1101/2021.06.24.449770

**Authors:** Luis E. Martinetti, Kelly E. Bonekamp, Dawn M. Autio, Hye-Hyun Kim, Shane R. Crandall

## Abstract

Short-term plasticity regulates the strength of central synapses as a function of previous activity. In the neocortex, direct synaptic interactions between areas play a central role in cognitive function, but the activity-dependent regulation of these long-range corticocortical connections and their impact on a postsynaptic target neuron is unclear. Here, we use an optogenetic strategy to study the connections between mouse primary somatosensory and motor cortex. We found that short-term facilitation was strong in both corticocortical synapses, resulting in far more sustained responses than local intra-cortical and thalamocortical connections. A major difference between pathways was that the synaptic strength and magnitude of facilitation were distinct for individual excitatory cells located across all cortical layers and specific subtypes of GABAergic neurons. Facilitation was dependent on the presynaptic calcium sensor synaptotagmin-7 and altered by several optogenetic approaches. Current-clamp recordings revealed that during repetitive activation, the short-term dynamics of corticocortical synapses enhanced the excitability of layer 2/3 pyramidal neurons, increasing the probability of spiking with activity. Furthermore, the properties of the connections linking primary with secondary somatosensory cortex resemble those between somatosensory-motor areas. These short-term changes in transmission properties suggest long-range corticocortical synapses are specialized for conveying information over relatively extended periods.

## INTRODUCTION

The strength of a synapse depends on its recent history of activity, and the rules governing these use-dependent changes can vary widely from one synapse to another (Zucker and Regehr 2002). For example, repetitive activation leads to increased transmitter release (facilitation) from some synapses and decreases from others (depression), while many synapses respond with neither or both. These diverse forms of synaptic plasticity lend flexibility to neural circuits (Abbott and Regehr 2004), allowing communication to be regulated dynamically and selectively.

Communication within the neocortex is essential for sensation, motor control, and cognitive function. It contains three general excitatory connections: local intracortical synapses and long-range synapses from other cortical areas and the thalamus. Although the properties of thalamocortical and local cortical synapses are relatively well understood (Gil et al. 1999; Reyes and Sakmann 1999; Hempel et al. 2000; Feldmeyer et al. 2002; Bruno and Sakmann 2006), the effect of a signal transmitted from one cortical area to another is less clear. Obviously, understanding the properties of these synapses is essential to understanding corticocortical (CC) information processing and could provide insight into cortex-dependent processes such as attention, prediction, and awareness (Engel et al. 2001; Bastos et al. 2012; Gilbert and Li 2013).

Addressing this critical issue has been technically challenging, mainly because long-range CC synaptic pathways are not amenable to study using conventional electrophysiological approaches. For example, CC projections can be diffuse and reciprocal (Veinante and Deschenes 2003), making it difficult to access and accurately measure their synapses’ relative contribution *in vivo*. Furthermore, in isolated brain preparations, electrical stimulation of CC inputs is problematic due to the proximity of these axons to non-targeted cells/axons (Dong et al. 2004; Covic and Sherman 2011; Rocco-Donovan et al. 2011; Petrof et al. 2015).

Optogenetic strategies overcome many of these problems (Petreanu et al. 2007; Cruikshank et al. 2010; Collins et al. 2018), but there are several difficulties in using these approaches to study synapses (Jackman et al. 2014). In most studies using opsins, the emphasis has been on isolating monosynaptic CC responses during perfusion of sodium and potassium channel blockers (Petreanu et al. 2009; Mao et al. 2011; Hooks et al. 2013; Yang et al. 2013; D’Souza et al. 2016; Young et al. 2021). This approach helps determine anatomical connections, but it can distort short-term dynamics (Cruikshank et al. 2010). In other studies, the characterization of synapses was limited (Kinnischtzke et al. 2014; Petrof *et al*. 2015; Zolnik et al. 2020; Naskar et al. 2021), and they did not address the pitfalls associated with using opsins to study synapses, such as artificial depression (Zhang and Oertner 2007; Jackman et al. 2014).

Here we apply an optogenetic strategy that overcomes these difficulties to investigate the connections between mouse vibrissal primary sensory (vS1) and motor cortex (vM1) (White and DeAmicis 1977; Porter and White 1983), two areas essential for active sensation, motor execution, and sensorimotor integration (Kleinfeld et al. 2006; Diamond et al. 2008). Contrasting with most excitatory synapses in the neocortex, we find that monosynaptic CC responses underwent short-term facilitation and that this synaptic adjustment plays a critical role in controlling the excitability of pyramidal cells. Therefore, long-range CC synapses appear specialized for carrying signals over sustained periods.

## RESULTS

### Optical control of CC synaptic transmission

Here we applied optogenetic control strategies to examine the connections between vS1 and vM1 (Crandall et al. 2015; Crandall et al. 2017). We identified vM1 and vS1 using viral-mediated anterograde tracing and the presence of large L4 barrels, respectively (Figure S1). vM1 was defined as a narrow region in the posteromedial part of the frontal cortex that reciprocally connected with vS1, consistent with previous studies (Hoffer et al. 2003; Ferezou et al. 2007; Mao et al. 2011). Next, we injected an adeno-associated virus (AAV2) carrying genes for a channelrhodopsin-2/enhanced yellow fluorescent protein construct (ChR2-EYFP) in vS1 or vM1 of mice *in vivo*. Three weeks after vS1 injections, there was robust ChR2-EFYP expression in vS1 axons/terminals within vM1 (Figures 1A and S1A-D). vS1 terminal arbors formed a narrow band ascending from the white matter within vM1 and concentrated in layer 1 (L1) and L2/3, consistent with a previous report (Mao et al. 2011). Similarly, three weeks after vM1 injections, there was intense ChR2-EYFP expression in vM1 axons/terminals within vS1 (Figures 1B and S1E). vM1 terminal arbors covered a large expanse of vS1 and concentrated in L1 and L5/6, precisely where they are known to terminate (Veinante and Deschenes 2003). Brief LED flashes directly over cortical cells located in the injection site evoked large inward currents with very short onset latencies (<0.35 ms) that were resistant to blocking fast glutamatergic and GABAergic transmission, confirming they expressed ChR2 (Figure S3A).

**Figure 1.**
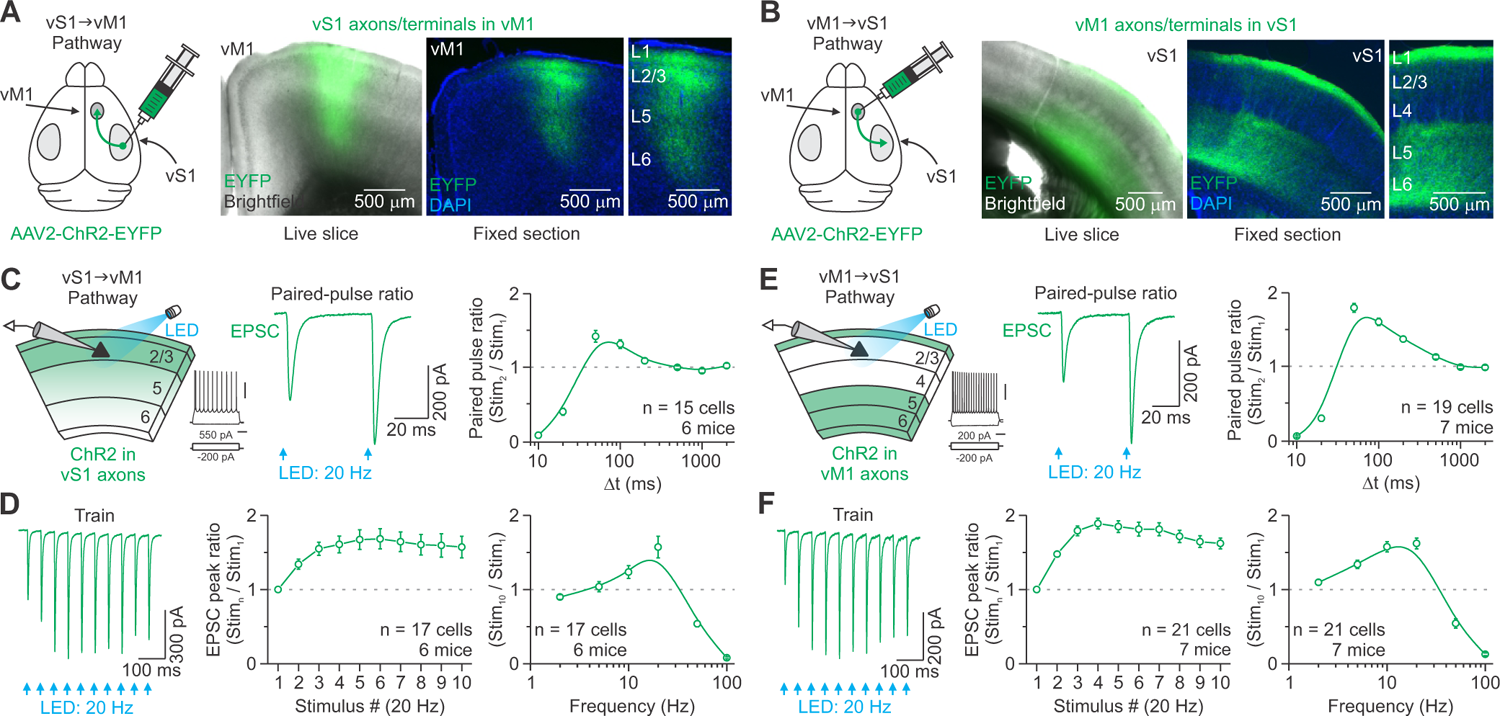
Optical stimulation of vS1 and vM1 CC projections evoked facilitating synaptic excitation in L2/3 neurons. (A-B) Left: Schematic showing virus injected into vS1 (A) or vM1 (B) of mice *in vivo*. Middle: Live slice (300 μm) image showing an overlay of EYFP with brightfield. Right: Low- and high-magnification fluorescence image of a 60-μm-thick section from the same live slice. EYFP-labeled vS1 axons terminate densely in superficial layers of vM1 (A), whereas EYFP-labeled vM1 axons terminate densely in L1 and L5/6 of vS1 (B). Sections were counterstained with DAPI. (C) Left: Recording schematic showing photostimulation of ChR2-expressing vS1 terminal arbors (green) and a whole-cell recording from an excitatory L2/3 cell in vM1. Responses of a non-expressing L2/3 RS cell to intracellular current steps (scale bars, 40 mV/200 ms). Middle: vS1-vM1 EPSCs evoked in the same neuron (shown in C, left) by a pair of optical stimuli at 20 Hz (blue arrow, 0.5 ms) (average of 10 trials). Right: Population data showing the average peak paired-pulse ratio at different interstimulus intervals (Δt). (D) Left: vS1-vM1 EPSCs evoked in the same cell (shown in C) by a 20 Hz train of optical stimuli (average of 10 trials). EPSCs increased 60-70% from the first to fourth pulse. Middle and Right: EPSC amplitudes plotted as a function of stimulus number within 20 Hz trains (normalized to first responses) and the peak responses to the tenth stimulus as a function of stimulus frequency (normalized to first responses). (E) Left: Recording schematic for the vM1-vS1 CC pathway. Responses of a non-expressing L2/3 RS cell to intracellular current steps (scale bars, 40 mV/200 ms). Middle: vM1-vS1 EPSCs evoked in the same neuron (shown in E, left) by a pair of optical stimuli at 20 Hz (blue arrows, 0.5 ms) (average of 16 trials). Right: Population data showing the average peak paired-pulse ratio at different interstimulus intervals (Δt). (F) Left: vM1-vS1 EPSCs evoked in the same neuron (shown in E) by a 20 Hz train of optical stimuli (average of 17 trials). EPSCs nearly doubled from the first to fourth pulse. Middle and Right: EPSC amplitudes plotted as a function of stimulus number within 20 Hz trains (normalized to first responses) and the peak responses to the tenth stimulus as a function of stimulus frequency (normalized to first responses). The light intensity for each cell was set to obtain an initial 200 pA EPSC when held near the inhibitory reversal potential (−94 mV, see Methods). Values are represented as mean ± SEM. See also Figures S1-S5 and Table S1.

Previous studies suggest that ChR2 desensitization impairs axons’ ability to follow during repetitive stimulation (Olsen et al. 2012; Hass and Glickfeld 2016), resulting in artificial synaptic depression. To determine the efficacy of ChR2 for activating CC pathways, we stimulated vM1 and vS1 axons/terminals with 20 Hz optical trains (10 pulses, 0.5 ms flashes, 1500 μm diameter spot) and measured effects on the extracellular, pharmacologically isolated fiber volley (Hass and Glickfeld 2016) (Figure S2). Fiber volley amplitude directly relates to the number of axons recruited by stimulation (Andersen et al. 1977). We found that optical stimulation evoked remarkably consistent fiber volley waveforms with modest amplitude decay (8.2 ± 3.8%) and minimal latency shift within trains (0.35 ± 0.05 ms; Figure S2E-G). Thus, 20 Hz optical stimulation of CC pathways is reliable under our conditions and can be used to study these synaptic connections.

### Long-range CC synapses express short-term facilitation

We first examined the pathway linking vS1 to vM1. We used voltage-clamp recordings to measure optically evoked excitatory postsynaptic currents (EPSCs) from excitatory regular-spiking (RS) neurons in L2/3 of vM1 (Figure 1C). Here the RS cells were identified using previously described physiological criteria (see Methods). We used brief, low-intensity LED stimuli to avoid mixed mono- and polysynaptic excitatory activity, keeping the initial synaptic current ∼200 pA when measured near the inhibitory reversal potential (−94 mV; see Methods). This intensity evoked subthreshold potentials for all cells tested (∼3 mV) (Table S1).

For all L2/3 RS cells recorded near ChR2-expressing vS1 arbors within vM1, brief LED flashes evoked fast EPSCs with onset latencies averaging 2.34 ± 0.04 ms (Figure S3A). These onsets are consistent with the timing of neurotransmission at fast synapses (Sabatini and Regehr 1999) and indicate that cells did not express ChR2. We next used paired-pulse stimulation to assess short-term plasticity (Zucker and Regehr 2002). Pairs of closely spaced optical flashes resulted in facilitation for interstimulus intervals of 50-200 ms and a peak enhancement of ∼1.5-fold (Figure 1C). However, we observed intense short-term depression for interstimulus intervals of 10-20 ms, consistent with reports that ChR2 does not reliably evoke spikes above 25 Hz (Lin et al. 2009). Since presynaptic neurons often discharge several action potentials *in vivo*, we also assessed synaptic dynamics using trains of flashes. Consistent with paired-pulse stimulation, all responses underwent facilitation during 5-20 Hz optical trains (Figure 1D and S3D).

We next examined the reciprocal pathway linking vM1 to vS1 (Figure 1E). Again we recorded from L2/3 RS cells, which receive direct input from vM1 on their apical dendrites in L1 (Petreanu et al. 2009). For most cells located near ChR2-expressing vM1 axons/terminals in L1, brief photostimulation evoked clear synaptic responses with short onset latencies (1.79 ± 0.03 ms; Figure S3A). When we used paired-pulse stimulation, cells responded with an increase in synaptic strength for short interstimulus intervals of 50-500 ms and a peak enhancement of ∼2-fold (Figure 1E). Furthermore, all responses evoked by 2-20 Hz optical trains exhibited facilitation (Figure 1F and S3E).

The short onset latencies of the CC responses suggest they are likely mediated by fast ionotropic glutamate receptors. Indeed, the combined application of the selective NMDA and AMPA kainate-type glutamate receptor antagonists APV and DNQX (50 and 20 μM, respectively) eliminated the responses in all cells tested (6/6 cells from 4 mice) (Figure S3B). Responses were also abolished after blocking sodium channels with tetrodotoxin (TTX: 1 μM) but returned with the subsequent addition of the potassium channel blocker 4-aminopyridine (4AP: 1 mM; 5/5 cells from 2 mice) (Figure S3C). The latter finding confirms that the light-evoked responses were both action potential-dependent and monosynaptic (Petreanu et al. 2009; Cruikshank et al. 2010).

To rule out the possibility that the observed facilitation reflects the superposition of long-range and local inputs, we compared the responses evoked under control conditions with those recorded on trials that we reduced the likelihood of postsynaptic firing (Figure S4). We suppressed the responsiveness of local excitatory cells either optogenetically by activating cortical parvalbumin (PV)-expressing interneurons that conditionally expressed the red-shifted opsin ChrimsonR (Klapoetke et al. 2014) or pharmacologically by blocking nonlinearities in the synaptic conductances associated with NMDA receptors (APV: 50 μM) (Larkum et al. 2009). We found that silencing local excitatory cells during the activation of long-range projections did not affect the observed facilitation (Figure S4), indicating that local inputs do not confound the CC responses.

To determine if our light stimulation method, which includes over-terminal illumination, significantly blunted short-term facilitation, we compared responses evoked by our approach and over-axon stimulation, which produces more physiological responses (Jackman et al. 2014). We found that the two stimulation methods produced similar dynamics during 20 Hz trains (Figure S5), suggesting that our stimulation method offers a reasonable strategy for assessing the short-term plasticity of CC synapses.

Altogether, these results established that reciprocal CC connections between vS1 and vM1 give rise to facilitating synapses capable of fast sustained excitatory transmission during brief periods of elevated presynaptic activity, contrasting with several previous reports (Lee et al. 2013; Kinnischtzke *et al*. 2014; Petrof et al. 2015; Naskar et al. 2021).

### The temporal dynamics of long-range and local intracortical excitation

The finding that long-range CC responses underwent short-term facilitation is somewhat surprising and intriguing given a key feature of most local excitatory–to– excitatory connections within the cortex is short-term depression (Gil et al. 1999; Reyes and Sakmann 1999; Feldmeyer et al. 2002). To better understand how different excitatory–to–excitatory connections within the neocortex are differentially affected by activity, we compared evoked responses at three general classes of excitatory synapses. At long-range CC synapses, 20 Hz optical stimulation resulted in paired-pulse facilitation (vS1-vM1 L2/3: 1.42 ± 0.08; vM1-vS1 L2/3: 1.80 ± 0.06; Figure 2A).

**Figure 2.**
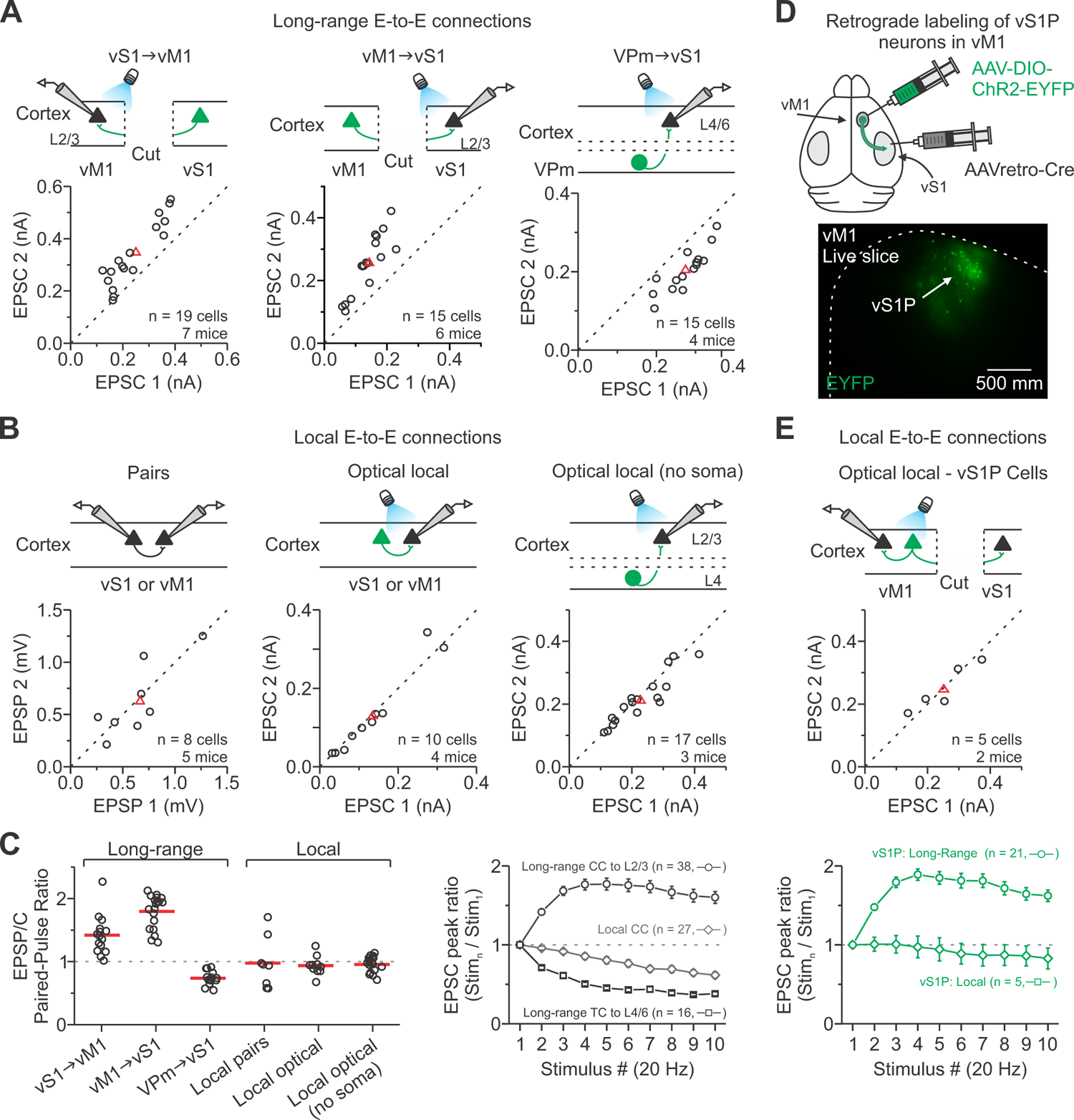
Synaptic responses during repetitive activation are more sustained for long-range CC than other excitatory cortical connections. (A) Long-range excitatory-to-excitatory (E-to-E) connections. A comparison of the first and second EPSC amplitude evoked by a pair of optical stimuli (20 Hz) directed at long-range CC or TC axons in the cortex: vM1-vS1_L2/3_, vS1-vM1_L2/3_, and VPm-vS1_L4/6_. (B) Local E-to-E connections. A comparison of the first and second EPSP/C amplitude evoked by a pair of stimuli for local excitatory connections in the cortex: pairs of excitatory cells (20-60 Hz), photostimulation of local excitatory neurons/axons that conditionally expressed ChR2 in *Rbp4-Cre* or *Scnn1a-Cre-Tg3* mice (10-20 Hz), and photostimulation of L4 to L2/3 axons/terminals (without parent somata) that conditionally expressed ChR2 using *Scnn1a-Cre-Tg3* mice (20 Hz). To isolate axons/terminals from parent somata, a cut was made between L4 and L2/3. (C) Left: Summary graph of paired-pulse ratios for the long-range and local excitatory cortical connections. Paired-pulse ratios were significantly larger for both long-range CC connections than all other excitatory connections (p < 0.0001, One-way ANOVA with Bonferroni’s post-hoc test). Right: Summary graph shows EPSC amplitudes plotted as a function of stimulus number within 20 Hz trains. Synaptic dynamics were significantly different, with responses to long-range CC inputs exhibiting short-term facilitation (n = 38 cells, 13 mice), local connections displaying weak depression (n = 27 cells, 7 mice), and long-range thalamocortical inputs showing strong depression (n = 16 cells, 4 mice) (p < 0.00001, Two-way ANOVA, stim. 2-10). (D) Top: Schematic showing AAVretro-Cre injected into the vS1 and Cre-dependent AAV-DIO-ChR2 injected into ipsilateral vM1 of mice *in vivo*. Bottom: Live slice (300 μm) image of vM1 showing EYFP fluorescence indicating the location of ChR2-EYFP. (E) Top: A comparison of the first and second EPSC amplitude evoked by a pair of optical stimuli (20 Hz) directed at ChR2-expressing vS1P cells in vM1 (p = 0.93, Paired t-test). Bottom: Summary graph shows EPSC amplitudes plotted as a function of stimulus number within 20 Hz trains for local and long-range targets of vS1P cells in vM1 (Local: n = 5 cells, 2 mice; Long-range: 21 cells, 7 mice; p < 0.00001, Two-way ANOVA, stim. 2-10). Red triangles and lines represent means. Long-range CC data from Figure 1. Values are represented as mean ± SEM.

This facilitation contrasts sharply with long-range subcortical inputs to the cortex from the thalamus, such as core sensory thalamocortical synapses (VPm-vS1 L4/L6), that exhibited robust paired-pulse depression when tested under identical conditions (0.71 ± 0.02; Figure 2A) (Gabernet et al. 2005; Cruikshank et al. 2010). Conversely, local connections between pairs of excitatory cells in vS1 or vM1 were more stable at 20 Hz, depressing only slightly across the tested population (0.98 ± 0.14; Figure 2B). Optical stimulation of local ChR2-expressing cortical cells (0.94 ± 0.05) or axons/terminals without parent somata evoked similar paired-pulse ratios (0.95 ± 0.05; Figure 2B).

These data demonstrate that the synaptic dynamics of three common types of glutamatergic synapses in the neocortex are dramatically different (p < 0.0001, Two-way ANOVA, stim. 2-10; Figure 2C), suggesting that they might serve dynamically distinct roles in neocortical operations.

Remarkably, local and long-range CC input to excitatory cells displayed different synaptic properties, suggesting that local and long-distance synaptic terminals of the same pyramidal neurons could exhibit different release properties. To test this hypothesis, we next injected AAVretro-Cre into vS1 to gain retrograde access to vS1-projecting(P) neurons in vM1 and then a Cre-dependent AAV-DIO-ChR2-EYFP into the ipsilateral vM1 to optically control the activity of these cells (Figure 2D) (Tervo et al. 2016). After allowing for expression, we observed ChR2-EYFP expression in vS1P neurons in vM1. We next recorded non-expressing cells in vM1 to study the dynamics of vS1P inputs onto local excitatory neurons. In contrast to our finding for long-range vM1- vS1 synapses, we found that local vS1P-evoked EPSCs depressed (p < 0.0001, Two-way ANOVA, stim. 2-10; Figure 2E). These findings imply that the presynaptic properties of CC projection neurons can differ according to whether the target cortical neuron is in their local circuit or a distant cortical area.

### Short-term plasticity of CC input as a function of cortical layer

Given the stark laminar differences in the spatial arrangement of vS1 and vM1 projections (Figure 1), and previous details about long-range CC connectivity (Petreanu et al. 2009; Mao et al. 2011; Kinnischtzke et al. 2014), we wondered if the short-term dynamics of these synapses depended on cortical layer. To investigate this, we compared the responses in excitatory cells located across the vertical depth of the cortex using layer-specific Cre-recombinase driver mouse lines crossed with a tdTomato (tdT) reporter (see Methods and Table S2). To control for variability in ChR2 expression in different slices and mice, we sequentially recorded from an identified L2/3 RS cell and an excitatory neuron (tdT-positive and negative) in a separate layer of the same column.

In vM1, excitatory vS1 currents were strongest in L2/3 and increased 50-70% during 20 Hz trains, whereas currents in L5/6 cells were significantly weaker, initially facilitated (10-15%), and then depressed with repetitive stimulation (Figure 3A-C). In contrast, in vS1, excitatory vM1 currents were strongest in L6, and responses approximately doubled during 20 Hz trains in L2/3 and L6, whereas short-term plasticity was highly heterogeneous across L5 connections (Figure 3D-F). Together, these results indicate that many long-range CC synapses undergo short-term facilitation, with absolute strength and dynamics differing with layer.

**Figure 3.**
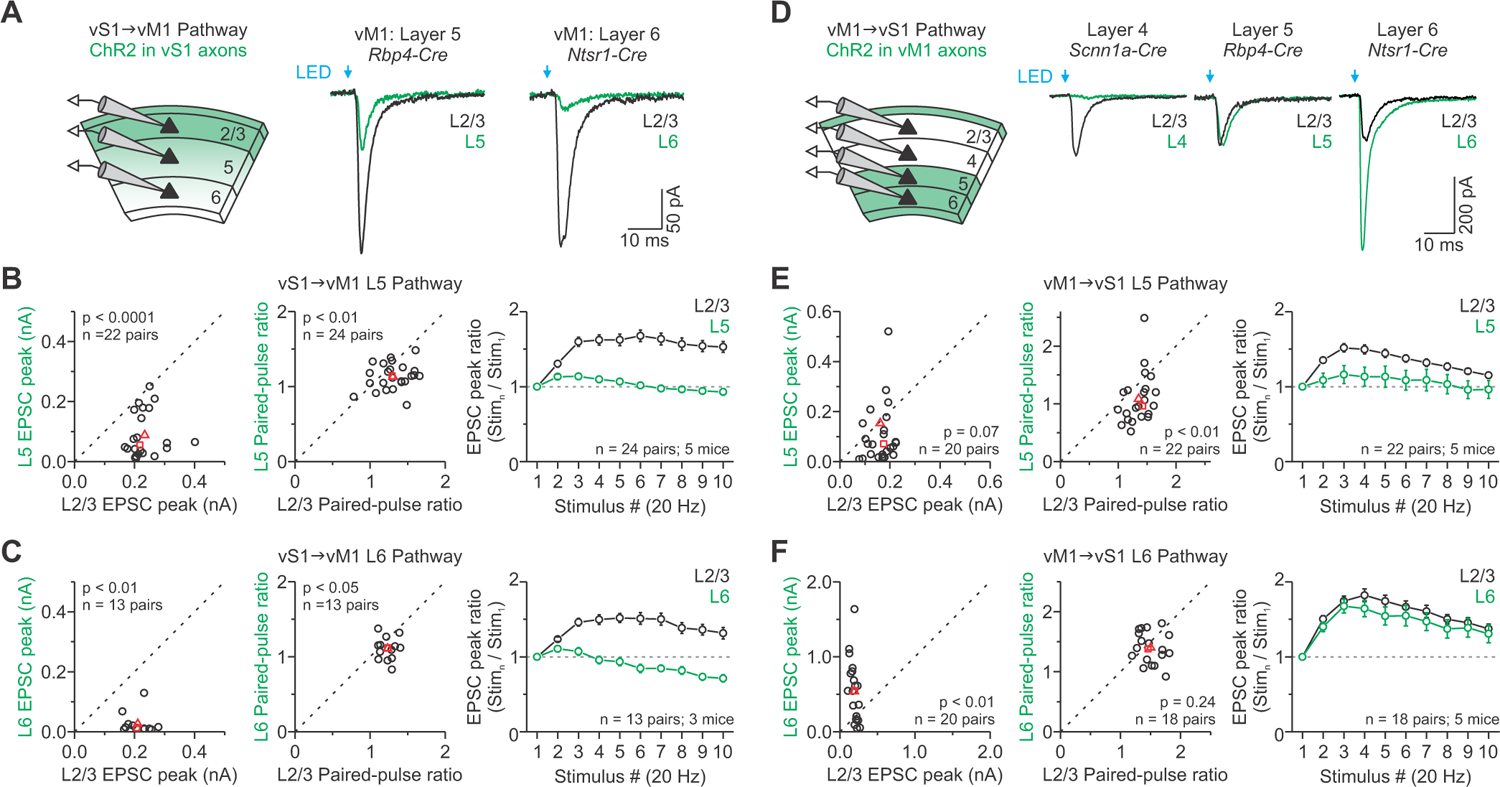
Comparison of ChR2-evoked CC responses across layers. (A) Left: Recording schematic for the vS1-vM1 pathway. We sequentially recorded an excitatory L2/3 cell and an excitatory neuron in a separate layer of the same column. Layers were determined using layer-specific Cre-driver mouse lines crossed with a tdTomato reporter (see Methods). Right: Representative single EPSCs evoked optically for pairs of excitatory cells recorded in different layers of vM1 (average of 6-11 trials). (B-C) A comparison of the average vS1-vM1 optically evoked EPSC amplitude (left), paired-pulse ratio at 20 Hz (middle), and EPSC amplitudes plotted as a function of stimulus number within 20 Hz trains for each L2/3-L5 (B) and L2/3-L6 cell pair (C). Red triangles and squares represent means and median, respectively. n displayed on the plot. L5 and L6 responses were significantly lower (p < 0.01, Wilcoxon paired signed-rank test), displayed lower paired-pulse ratios (p < 0.0001, Paired t-test), and underwent less facilitation during 20 Hz trains than those in L2/3 (p < 0.0001, Two-way ANOVA, stim. 2-10). (D) Left: The same experimental approach described in (A) for the vM1-vS1 pathway. Right: Representative single EPSCs evoked for pairs of excitatory cells in different layers of vS1 (average of 5-11 trials). (E-F) A comparison of the average vM1-vS1 optically evoked EPSC amplitude (left), paired-pulse ratio at 20 Hz (middle), and EPSC amplitudes plotted as a function of stimulus number within 20 Hz trains for each L2/3-L5 (E) and L2/3-L6 cell pair (F). Red triangles and squares represent means and median, respectively. n displayed on the plot. EPSCs were stronger in L6 (p < 0.01, Paired t-test), weaker in L4 (n = 7 pairs, 3 mice, data not shown, p < 0.03, Wilcoxon paired signed-rank test), and similar in L5 as compared to those in L2/3 (p = 0.07, Wilcoxon paired signed-rank test). L5 responses underwent less facilitation than those in L2/3 (PPR: p < 0.01, Mann-Whitney U test; Train: p < 0.0001, Two-way ANOVA, stim. 2-10), whereas L6 responses displayed similar paired-pulse facilitation (p = 0.24, Paired t-test) but facilitated less than L2/3 during 20 Hz trains (p < 0.01, Two-way ANOVA, stim. 2-10). L4 responses were too weak to test dynamics. For single EPSCs, cells were tested at the light intensity needed to obtain an initial 200 pA EPSC in L2/3 in voltage-clamp at −94 mV, whereas short-term plasticity was tested at the cell’s own 200 pA intensity. Values are represented as mean ± SEM. See also Figure S2 and Tables S1-S2.

### Properties of excitatory CC synapses onto different L2/3 interneuron subtypes

Long-range CC connections in vS1 and other cortical areas also target GABAergic inhibitory interneurons (Dong et al. 2004; Lee et al. 2013; Yang et al. 2013; Kinnischtzke et al. 2014; Zhang et al. 2014; Naskar et al. 2021). To compare CC responses among different subtypes of L2/3 interneurons, we measured optically evoked CC responses from an identified interneuron and a neighboring RS excitatory cell. To target different L2/3 interneuron subtypes, we crossed three well-established Cre-driver mouse lines with tdT reporter mice (Ai14) to label PV-, somatostatin (SOM)-, or vasoactive intestinal peptide (VIP)-expressing interneurons (see Methods).

Figures 4A and 4E show optically evoked CC responses from an interneuron and neighboring RS cell. We found that EPSCs recorded from PV cells had the largest amplitudes and showed slightly less facilitation at 20 Hz than RS cells for both pathways (EPSC peak normalized to RS: vM1 PV cells, 13.3 ± 5.8, median = 4.9; vS1 PV cells, 6.1 ± 1.4, median = 4.8; Figure 4B, F). EPSCs from SOM cells had much smaller amplitudes for both CC pathways but displayed dramatic short-term facilitation at 20 Hz (EPSC peak normalized to RS: vM1 SOM cells, 0.17 ± 0.03, median = 0.18; vS1 SOM cells, 0.66 ± 0.11, median = 0.55; Figure 4C, G). In contrast to those of PV and SOM cells, the EPSCs of VIP cells were significantly weaker in the vS1-vM1 pathway, whereas the EPSC amplitudes were statistically similar to RS cells in the vM1-vS1 pathway (EPSC peak normalized to RS: vM1 VIP cells, 0.14 ± 0.06, median = 0.10; vS1 VIP cells, 0.89 ± 0.28, median = 0.64; Figure 4D, H). Furthermore, the excitatory currents in VIP cells initially facilitated (10-45%) and then depressed with repetitive stimulation for both pathways (Figure 4D, H). Despite similar EPSC amplitudes, vM1 stimuli always evoked action potentials in the VIP cell (8/8 pairs) but rarely in the RS cell (1/8 pairs) (Figure S6). Altogether, these data suggest that long-range CC synapse strength and dynamics depend on the postsynaptic interneuron subtype and pathway.

**Figure 4.**
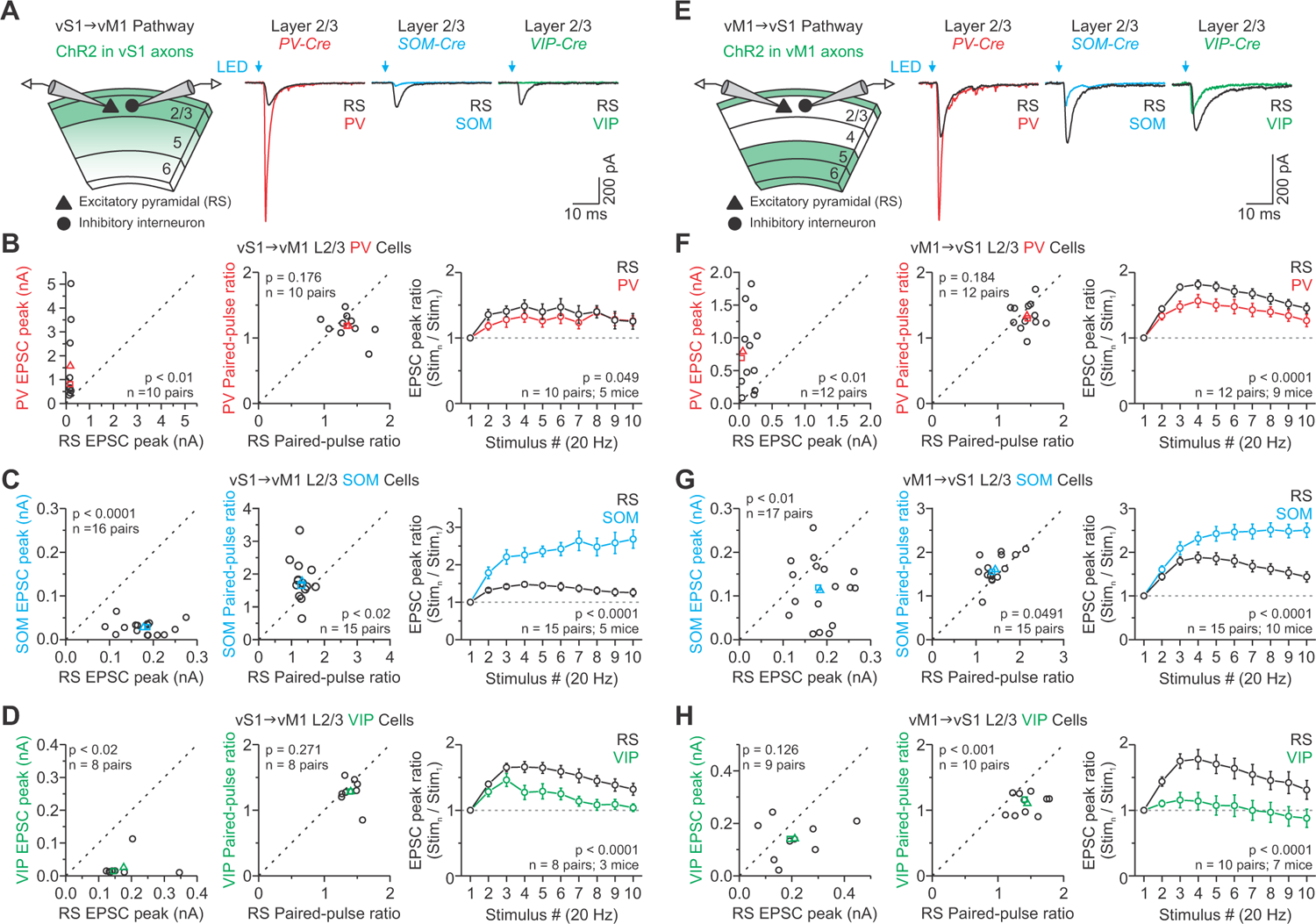
Comparison of ChR2-evoked CC responses across GABAergic interneurons in L2/3 of vM1 and vS1. (A) Left: Recording schematic for the vS1-vM1 pathway. We simultaneously recorded a specific L2/3 interneuron and a nearby excitatory neuron. Experiments utilized Cre-driver mice targeting three classes of GABAergic cells in the neocortex (see Methods). Right: Representative single EPSCs evoked optically in interneuron-RS pairs (average of 7-10 trials). (B-D) A comparison of the average vS1-vM1 optically evoked EPSC amplitude (left), the paired-pulse ratio at 20 Hz (middle), and EPSC amplitudes plotted as a function of stimulus number within 20 Hz trains for PV-RS (B), SOM-RS (C), and VIP-RS pairs (D). Colored triangles and squares represent means and median, respectively. n displayed on the plot. EPSCs were stronger in PV (p < 0.01, Wilcoxon paired signed-rank test) but weaker in both SOM and VIP as compared to those in L2/3 RS cells (SOM: p < 0.0001, Paired t-test; VIP: p < 0.02, Wilcoxon paired signed-rank test). Short-term dynamics of excitatory vS1-vM1 synapses onto L2/3 interneurons were significantly different during 20 Hz trains than RS cells (p < 0.05, Two-way ANOVA, stim. 2-10). (E) Left: The same experimental approach described in (A) for the vM1-vS1 pathway. Right: Representative single EPSCs evoked optically in interneuron-RS pairs (average of 8-10 trials). (F-H) A comparison of the average vM1-vS1 optically evoked EPSC amplitude (left), the paired-pulse ratio at 20 Hz (middle), and EPSC amplitudes plotted as a function of stimulus number within 20 Hz trains for PV-RS (F), SOM-RS (G), and VIP-RS pairs (H). Colored triangles and squares represent means and median, respectively. n displayed on the plot. EPSCs were stronger in PV (p < 0.01, Paired t-test), weaker in SOM (p < 0.01, Paired t-test), and similar in VIP as compared to those in L2/3 RS cells (p = 0.126, Paired t-test). Short-term dynamics of excitatory vS1-vM1 synapses onto L2/3 interneurons were significantly different during 20 Hz trains than RS cells (p < 0.0001, Two-way ANOVA, stim. 2-10). For single EPSCs, we tested pairs at the light intensity needed to obtain an initial 200 pA EPSC in the L2/3 excitatory neuron when recorded in voltage-clamp at −94 mV. We tested short-term plasticity at an intensity that evoked a reliable EPSC (typically 50-300 pA) for each cell. Values are represented as mean ± SEM. See also Figure S6.

### Experimental strategies that affect CC responses

Our results establish that facilitation is the predominant communication mode for conveying information between vS1 and vM1. To understand why our data differ from those previously reported, we first compared the effect of AAV serotype on optically evoked short-term synaptic plasticity (Jackman et al. 2014). Here, we injected mice with an identical vector to our previous work using AAV2 but with a different viral serotype (AAV1, AAV5, and AAV9; same titer). All three vectors produced robust ChR2-EYFP expression, similar to levels observed using AAV2 (Figure S7A). When we drove ChR2 expression by AAV5, responses evoked by optical trains exhibited less facilitation for the vS1-vM1 synapse but not for the vM1-vS1 synapse (Figure 5A and 5B). However, when using AAV1 to express ChR2, facilitation was weaker for the vM1-vS1 synapse but not for the vS1-vM1 synapse (Figure 5A and 5B). Responses were not different from those obtained using AAV2-ChR2 when using AAV9-ChR2. When we examined the effects of different fluorescent fusion proteins on optically evoked responses, we also found that currents evoked by ChR2-mCherry exhibited significantly less facilitation than ChR2-EYFP (Figure 5C), perhaps due to the overall poor surface expression observed with ChR2-mCherry (Figure S7B). These results indicate that using AAV1 and AAV5 produce only modest facilitation in a pathway-dependent manner and that mCherry tagging of ChR2 caused adverse effects.

**Figure 5.**
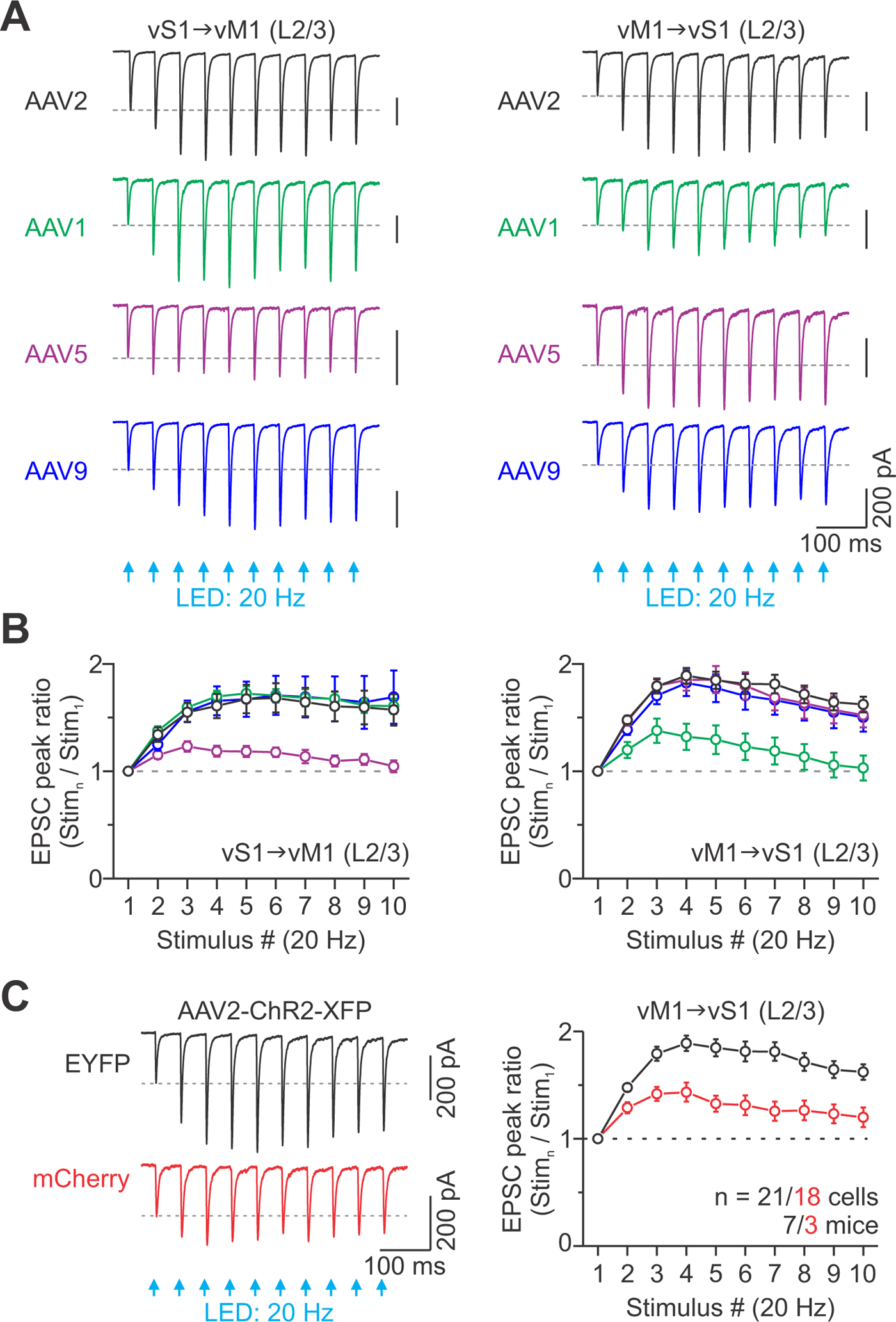
Effect of AAV serotype and ChR2 fusion protein on optically evoked facilitation at CC synapses. (A) For each serotype (AAV2, AAV1, AAV5, and AAV9), representative EPSCs evoked optically by 20 Hz trains for an excitatory L2/3 cell located in vM1 (left) or vS1 (right) (average of 3-17 trials). We used the same vector (AAV-hSyn-hChR2(H124R)-EFYP), titer (3.1 x 10^12^ viral genomes/ml), volume per injection site (∼0.15 μl), and transduction time (21±1 days) for each serotype to keep expression levels similar. (B) Summary graphs show the average EPSC amplitudes plotted as a function of stimulus number within 20 Hz trains for each serotype tested in the vS1-vM1 (left) and vM1-vS1 pathway (right). Facilitation was significantly weaker than other serotypes when using AAV5 in the vS1-vM1 pathway (p < 0.0001) and AAV1 in the vM1-vS1 pathway (p < 0.0001, Two-way ANOVA, stim. 2-10, with Bonferroni’s post-hoc test) (vS1-vM1: AAV1, n = 18 cells, 2 mice; AAV5, n = 16 cells, 3 mice; AAV9, n = 8 cells, 2 mice) (vM1-vS1: AAV1, n = 8 cells, 3 mice; AAV5, n = 8 cells, 3 mice; AAV9, n = 21 cells, 7 mice). AAV2 data same as shown in Figure 1D and 1F. (C) Left: Representative EPSCs evoked optically by 20 Hz trains for excitatory L2/3 cells in vS1 when using AAV2 to drive ChR2-EYFP or ChR2-mCherry expression in vM1 axons (average of 10 and 8 trials). For these experiments, we used the same titer and expression time. Right: Summary graph shows the average EPSC amplitudes plotted as a function of the stimulus number within 20 Hz trains for each vector. Short-term facilitation was blunted when using AAV2-ChR2-mCherry (n = 18 cells, 3 mice for mCherry; p < 0.0001, Two-way ANOVA, stim. 2-10). AAV2-ChR2-EYFP data same as shown in Figure 1F. Values are represented as mean ± SEM. See also Figures S7.

We also encountered several cells intermingled with ChR2-EYFP expressing axons/terminals that were light-sensitive when using AAV1, AAV5, and AAV9 (Figure S7C-E). However, we rarely encountered ChR2 expressing cells when using AAV2. These observations suggest that retrograde spread within the cortex is more frequent with some AAV serotypes.

Given direct excitation of ChR2-expressing terminals with light can cause artificial depression (Zhang and Oertner 2007; Jackman et al. 2014), we next wondered if the power density of over-terminal illumination affects the magnitude of facilitation at CC synapses. Our approach was to stimulate using brief wide-field illumination (1500 μm diameter spot) and low light power density to minimize complications that may arise due to direct stimulation of presynaptic terminals. To assess the impact of a more focused over-terminal illumination, we compared optical responses evoked by the same total light power but with light-spots of different diameters (1500 μm or 150 μm) (Figure 6A). We found that restricting the excitation light to the distal ends of ChR2-expressing axons caused the amplitude of facilitation to decrease significantly during repetitive stimulation (Figure 6B and 6C). We also found that pulse duration disrupted paired-pulse facilitation in a pathway-dependent manner (Figure S8A-B).

**Figure 6.**
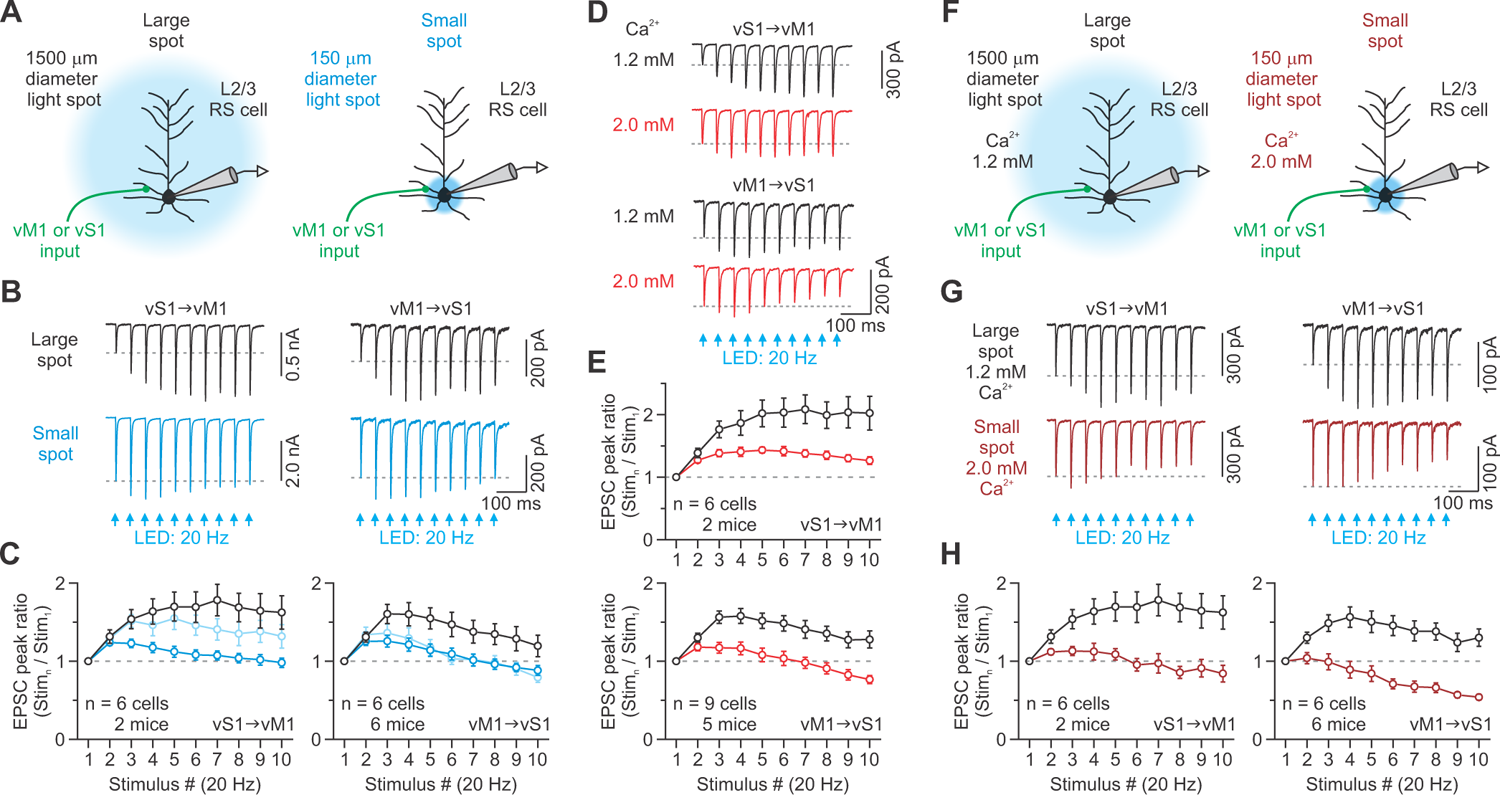
Effect of over-terminal stimulation and extracellular Ca^2+^ concentrations on optically evoked facilitation at CC synapses. (A) Recording schematic for (B-C) showing photostimulation of CC arbors using different light-spot sizes (blue circles). Restricting the excitation light to over the terminal ends of ChR2-expressing axons and keeping the total emitted light power the same will likely cause more terminal depolarization. (B) Representative EPSCs evoked by large- and small-field repetitive photostimulation at 20 Hz for an excitatory L2/3 cell in vM1 (left) or vS1 (right) (average of 5-11 trials). (C) Summary graphs show average EPSC amplitudes plotted as a function of stimulus number within 20 Hz trains for each condition. Restricting the excitation light significantly decreased the facilitation at both vS1-vM1 (p < 0.0001) and vM1-vS1 synapses (p < 0.0001, Two-way ANOVA, stim. 2-10; Average Power densities: vS1-vM1 = 0.71 and 71 mW/mm^2^; vS1-vM1 = 2.34 and 234 mW/mm^2^). Adjusting the power of the small spot to obtain an initial 200 pA EPSC also decreased facilitation (Light blue traces: vS1-vM1, p < 0.006; vM1-vS1, p < 0.0001; Two-way ANOVA, stim. 2-10; Average Power densities: vS1-vM1 = 21 mW/mm^2^; vS1-vM1 = 111 mW/mm^2^). (D) Representative EPSCs evoked optically by 20 Hz trains for an excitatory L2/3 cell located in vM1 (top) or vS1 (bottom) recorded in 1.2 and 2.0 mM external Ca^2+^ (average of 10-12 trials). (E) Summary graphs show average EPSC amplitudes plotted as a function of stimulus number within 20 Hz train for 1.2 and 2.0 mM external Ca^2+^. Raising Ca^2+^ significantly decreased facilitation at both vS1-vM1 (p < 0.0001) and vM1-vS1 synapses (p < 0.0001, Two-way ANOVA, stim. 2-10). (F) Recording schematic for (G-H) showing the same photostimulation approach described in (A). Large-field photostimulation was done in 1.2 mM external Ca^2+^, whereas small-field photostimulation was done in 2.0 mM external Ca^2+^. (G) Representative EPSCs evoked by large- and small-field repetitive photostimulation at 20 Hz recorded in 1.2 and 2.0 mM external Ca^2+^, respectively, for an excitatory L2/3 cell located in vM1 (left) or vS1 (right) (average of 10-12 trials). (H) Summary graphs show average EPSC amplitudes plotted as a function of stimulus number within 20 Hz train for the two conditions. Restricting the excitation light while in 2.0 mM external Ca^2+^ changed facilitation to depression (vS1-vM1, p < 0.0001; vM1-vS1, p < 0.0001, Two-way ANOVA, stim. 2-10). The light intensity was set to obtain an initial ∼200 pA EPSC for C (light blue), D-E, and F-G. Right: Values are represented as mean ± SEM. See also Figures S8.

We next examined the Ca^2+^-dependence of optically evoked EPSCs and facilitation. It is well established that Ca^2+^ ions play a crucial role in short-term plasticity (Zucker 1999), and thus far, we have conducted our studies in the presence of 1.2 mM Ca^2+^, which is around physiological conditions (Somjen 2002). However, many studies are performed in the presence of 2.0 mM external Ca^2+^. We found that increasing Ca^2+^ caused a significant increase in EPSC amplitude and reduced paired-pulse facilitation for both synapses (Figure S8C-D). To test further the impact of Ca^2+^, we measured how EPSC trains evoked with repetitive optical stimulation changed with external Ca^2+^. Increasing the concentration of Ca^2+^ to 2.0 mM significantly reduced the magnitude of facilitation observed during 20 Hz trains (Figure 6D and 5E).

Our findings indicate that both light stimulation technique and external Ca^2+^ levels can strongly affect the magnitude of facilitation at CC synapses, suggesting that combining these two experimental approaches could convert facilitating responses to depression. To test this hypothesis, we compared CC responses evoked by our standard approach to those triggered by focused stimulation of the distal ends of ChR2-expressing axons while in the presence of 2.0 mM Ca^2+^ (Figure 6F). Under control conditions, the responses underwent strong short-term facilitation, whereas subsequent changes to the light and external Ca^2+^ levels led to synaptic currents that depressed (Figure 6G and 6H).

Overall, these results reveal how different AAV serotypes, fluorescent fusion proteins, methods of light stimulation, and Ca^2+^ concentrations can influence the short-term synaptic plasticity of CC synapses and perhaps the reliability of synaptically driven circuit activity.

### Synaptotagmin 7 mediates facilitation at long-range CC synapses

Recent work has shown that the slow presynaptic Ca^2+^ sensor, synaptotagmin 7 (Syt7), plays critical roles in synaptic transmission, especially in short-term facilitation at many central synapses (Jackman et al. 2016). Therefore, we tested whether Syt7 could mediate facilitation at CC synapses. Immunohistochemistry revealed a Syt7 pattern in the hippocampus and thalamus similar to a previous report (Figure S9A) (Jackman et al. 2016). In the neocortex, we observed that Syt7 was present in vM1 and vS1 of wildtype (WT) mice but was absent in Syt7 knockout (KO) animals (Figure 7A and 7B). In WT mice, Syt7 labeling was prominent in L1, where CC synapses are dense and are known to contact the dendrites of pyramidal cells (Cauller et al. 1998; Petreanu et al. 2009; Mao et al. 2011), suggesting a possible expression of Syt7 in long-range CC terminals.

**Figure 7.**
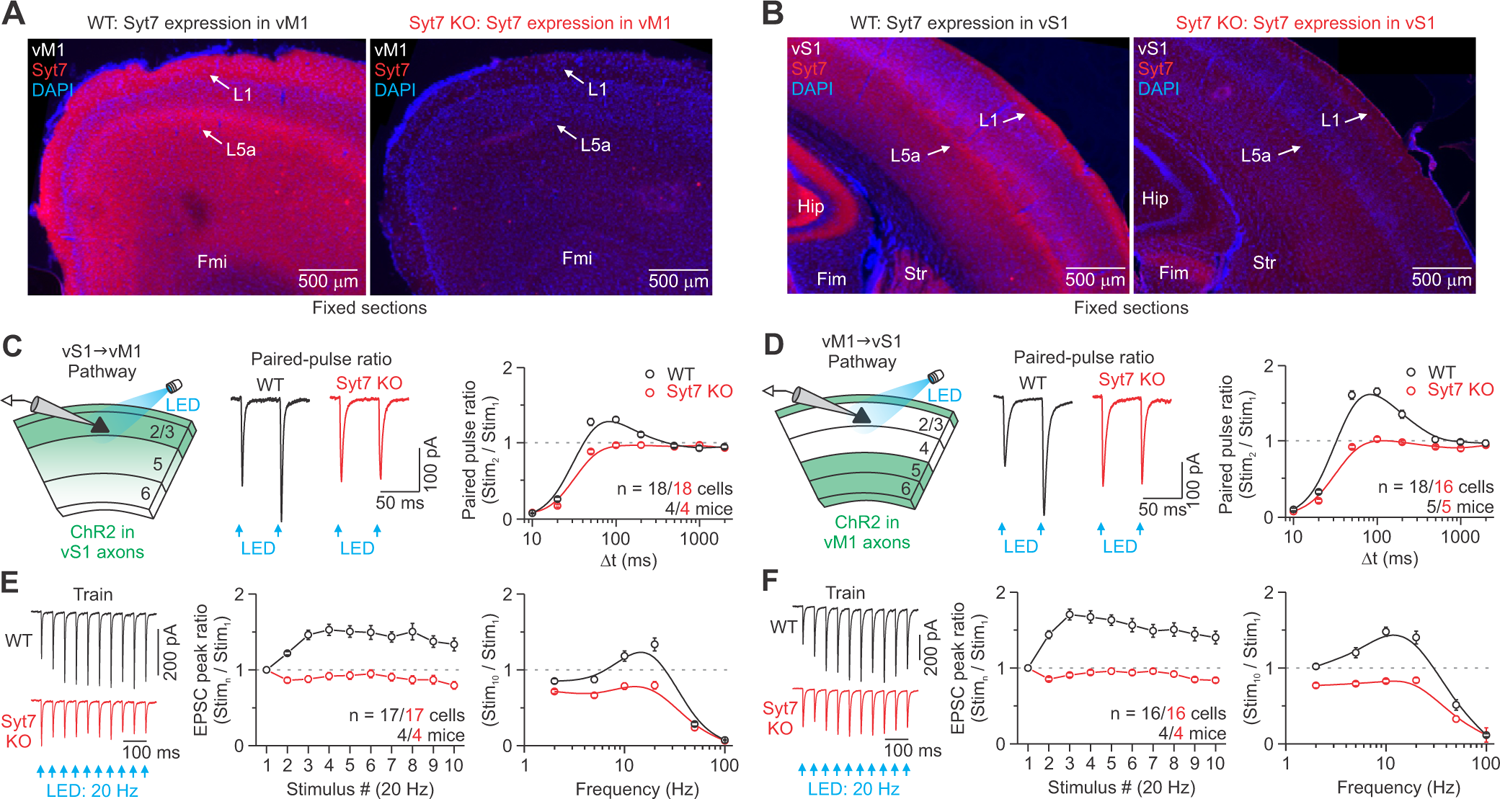
Facilitation at CC synapses is absent in Syt7 knockout mice. (A-B) Images of 60-μm-thick fixed coronal sections, centered on vM1 (A) and vS1 (B), from WT mice and Syt7 KO littermates (ages = P73 for A and P95 for B). Tissue was stained immunohistochemically for Syt7 and counterstained with DAPI. Syt7 expression was strong in L1 and L5a. Hip, hippocampus; Fim, fimbria of the hippocampus; Fmi, forceps minor of the corpus callosum, Str, striatum. (C-D) Left: Recording schematic for the vS1-vM1 (C) and vM1-vS1 CC pathway (D). The light intensity for each cell was set to obtain an initial 200 pA EPSC when held in voltage-clamp at −94 mV. Middle: Representative vS1-vM1 (C) and vM1-vS1 (D) EPSCs evoked in L2/3 RS cells by a pair of optical stimuli at 20 Hz, recorded from brain slices prepared from WT and Syt7 KO littermates (average of 9-17 trials). Right: Summary graphs show the average peak paired-pulse ratio at different interstimulus intervals (Δt) for vS1-vM1 (C) and vM1-vS1 synapses (D). n displayed on the plot. The peak paired-pulse ratio at 20 Hz was significantly different for WT and Syt7 KO at both synapses (p < 0.0001, Two-Sample t-test). (E-F) Left: vS1-vM1 (E) and vM1-vS1 (F) EPSCs evoked in the same pair of cells (shown in C and D) by a 20 Hz train of optical stimuli (average of 12-20 trials). Middle and Right: Summary graphs show EPSC amplitudes plotted as a function of stimulus number within 20 Hz trains, and the average normalized peak response to the tenth stimulus as a function of stimulus frequency for vS1-vM1 (E) and M1-vS1 synapses (F). In Syt7 KO mice, facilitation was eliminated at both synapses (p < 0.0001, Two-way ANOVA, stim. 2-10). Values are represented as mean ± SEM. See also Figures S9 and Table S3.

We next examined the functional role of Syt7 in facilitation at both CC synapses by performing similar optogenetic experiments in Syt7 KO mice and age-matched WT littermates. In WT mice, pairs of closely spaced optical stimuli and 5-20 Hz trains of flashes resulted in synaptic facilitation (Figure 7C-F). In Syt7 KO mice, facilitation was eliminated (Figure 7C-F and S9C). The loss of facilitation in Syt7 KO mice cannot be accounted for by the inability of ChR2 to drive presynaptic axons because optical stimulation evoked fiber volleys comparable to that of WT animals (Figure S9D-E). Nor can it be explained by differences in intrinsic properties between WT and KO cortical cells (Table S3). Altogether, these data indicate that Syt7 mediates short-term facilitation at CC synapses linking vS1 and vM1.

### Role of short-term dynamics in the direct modulation of L2/3 excitability

Having determined the short-term dynamics of CC synapses, we next examined how these inputs might control the excitability of L2/3 pyramidal neurons. The short-term plasticity of excitatory CC synapses suggests that they may influence L2/3 processing by directly increasing the overall excitation and spike probability of pyramidal neurons with repetitive activity. To test this idea, we recorded in current-clamp from L2/3 RS neurons and performed optical stimulation with 20 Hz trains (Figure 8A and 8B). During repetitive stimulation, the membrane potential of L2/3 vM1 and vS1 cells depolarized progressively, reaching its peak towards the ends of the trains. The overall magnitude of facilitation observed with the excitatory postsynaptic potentials (EPSPs) was consistent with the dynamics of the previously recorded EPSCs (Figure 8A and 8B). To determine the impact of CC input on L2/3 excitability, we next monitored the spiking behavior in response to 20 Hz trains while holding the cell at a depolarized membrane potential (−74 mV) (Figure 8C). Under these conditions, trains of optical stimuli reliably drove spiking in L2/3 RS cells. For both vM1 and vS1 neurons, the probability of spike discharge was lowest after the first stimulus and gradually increased following subsequent stimuli so that the maximal spiking probability occurred at the end of the train (Figure 8D). Together, these data indicate that direct CC effects on L2/3 excitability are regulated dynamically, mainly generating enhancement during brief periods of sustained activity.

**Figure 8.**
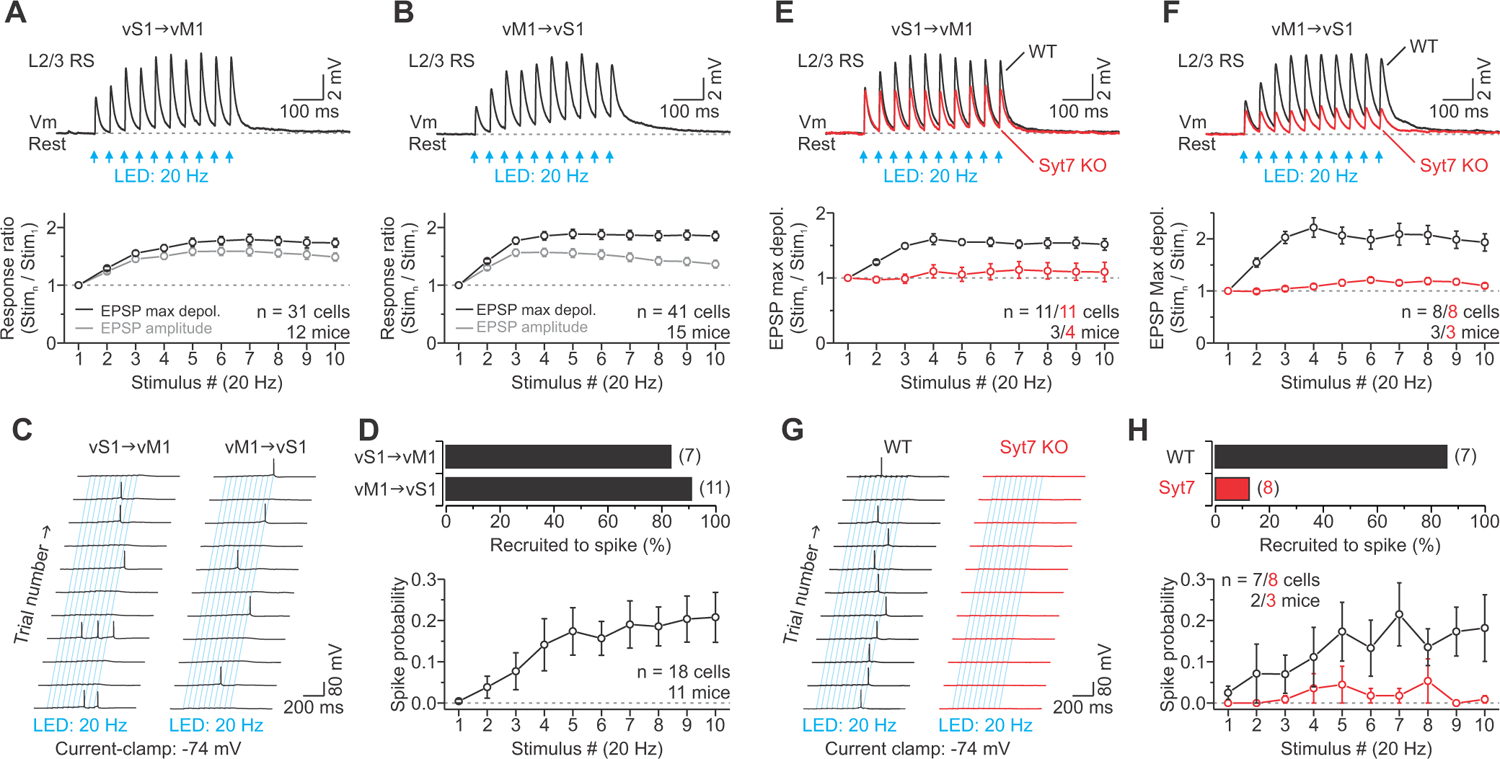
The functional role of short-term facilitation in the modulation of L2/3 excitability. (A-B) Top: Representative synaptic potentials evoked by low-intensity 20 Hz optical trains of CC arbors for an excitatory L2/3 cell at rest in vM1 (A) and vS1 (B) (average of 11 trials each). The light intensity for each cell was the same intensity needed to obtain an initial 200 pA EPSC when held in voltage-clamp at −94 mV. Bottom: Population data showing the average EPSPs’ maximal depolarization and EPSPs’ amplitude plotted as a function of stimulus number within 20 Hz trains for both vS1-vM1 (A) and vM1-vS1 (B) synapses. Short-term synaptic facilitation was apparent during 20 Hz trains. (C) Current-clamp recordings from an excitatory L2/3 cell in vM1 (left) and vS1 (right) in response to 20 Hz optical stimulation of CC axons/terminals. Same conditions as in (A-B), but neurons were held in current-clamp at a more depolarized membrane potential (−74 mV). A train of optical CC stimuli elicits action potentials in excitatory L2/3 cells in some but not all trials. (D) Top: Summary histogram showing the percentage of L2/3 RS cells recruited to spike by photostimulation of both CC pathways (n shown in parentheses). Bottom: Summary graph showing spike probability plotted against stimulus number for L2/3 RS cells in vM1 and vS1. (E-F) Top: Representative synaptic potentials evoked by low-intensity 20 Hz optical trains of CC arbors for an excitatory L2/3 neuron at rest in vM1 (A) and vS1 (B), recorded from brain slices prepared from WT and Syt7 KO mice (average of 11-12 trials). Bottom: Population data showing the average EPSPs’ maximal depolarization and EPSPs’ amplitude plotted as a function of stimulus number within the trains for both vS1-vM1 (E) and vM1-vS1 (F) synapses in WT and Syt7 KO mice. Short-term synaptic facilitation was absent in Syt7 KO mice (p < 0.0001, Two-way ANOVA, stim. 2-10). (G) Similar current-clamp recordings as described in (C) for excitatory L2/3 neurons recorded in brain tissue prepared from WT (left) and Syt7 KO (right) mice in response to 20 Hz optical stimulation of CC axons/terminals. A train of optical CC stimuli elicits action potentials in WT L2/3 cells in some but not all trials, whereas cells in KO mice rarely responded with action potentials. (H) Top: Summary histogram showing the percentage of L2/3 RS cells recruited to spike by photostimulation of CC afferents for both WT and Syt7 KO mice (n shown in parentheses). Bottom: Summary graph showing spike probability plotted against stimulus number for WT and Syt7 KO L2/3 RS cells. In Syt7 KO mice, spike probability was significantly reduced (p < 0.0001, Two-way ANOVA). Values are represented as mean ± SEM.

In general, the short-term synaptic dynamics of the CC evoked excitatory responses appeared to parallel the CC modulation of L2/3 excitability. To test the causal relationship between short-term facilitation and the changes in excitability, we performed similar current-clamp experiments in Syt7 KO and WT animals. In Syt7 KO mice, the facilitating CC-evoked EPSPs were abolished (Figure 8E and 8F). This loss of facilitating CC-evoked excitatory responses eliminated the late enhancement in L2/3 excitability (Figure 8G and 8H). These data indicate that the short-term dynamics of inputs to L2/3 cells can account for the CC-triggered late enhancement of their excitability.

### Long-range projections from vS1 to vS2 have similar synaptic properties

To determine whether other CC projections might share the same short-term plasticity described for the connections between vS1 and vM1, we examined the vibrissal secondary somatosensory cortex (vS2) and its inputs from vS1. We found that the short-term plasticity of vS1 synapses in L2/3 of vS2 was very similar to that of the vS1-vM1 CC system (Figure 9). These data suggest that short-term facilitation might be a general feature of long-range CC connectivity.

**Figure 9.**
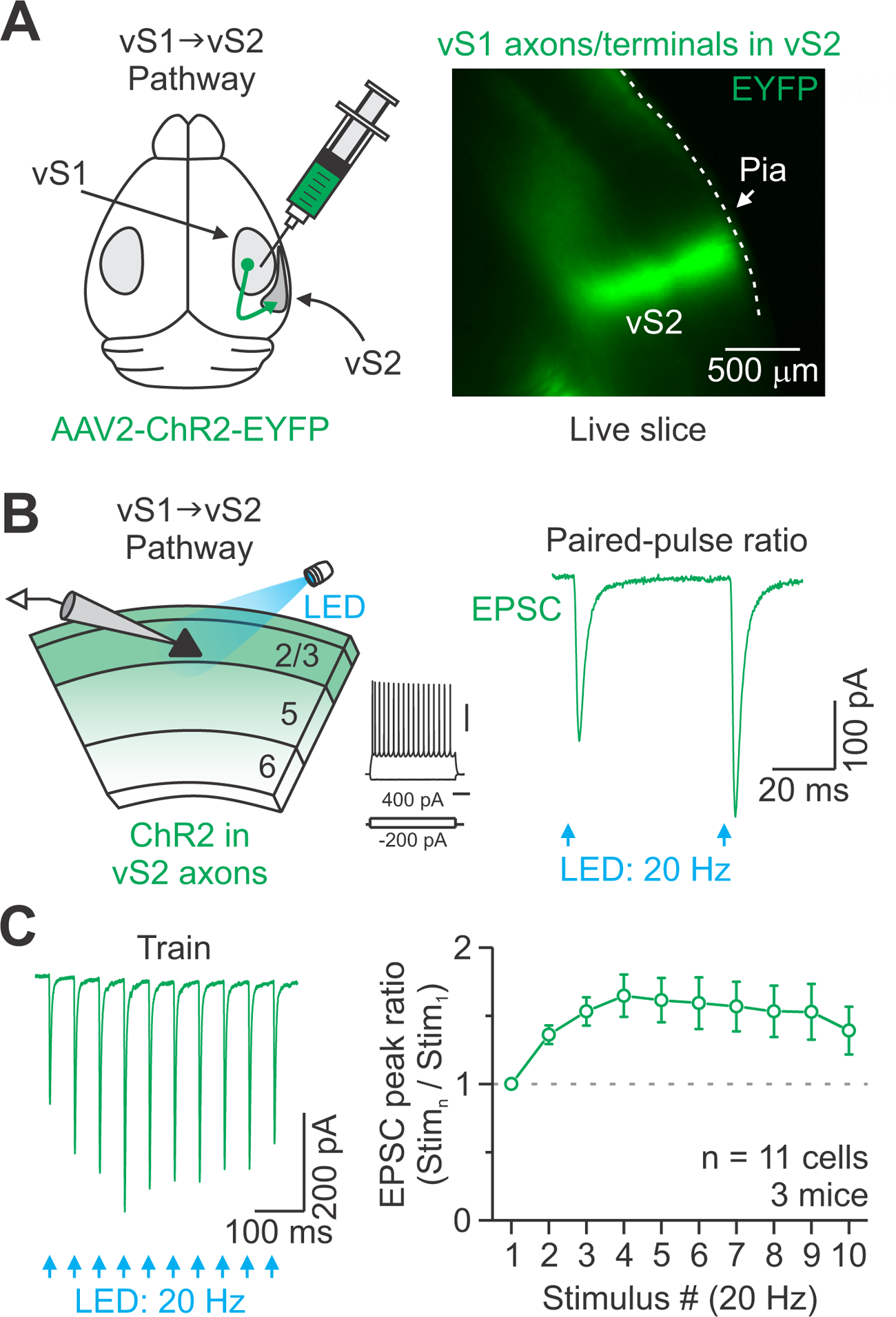
vS1 synapses in vS2 have similar dynamics as vS1-vM1 and vM1-vS1 synapses. (A) Right: Injection schematic. AAV2-ChR2-EFYP was injected unilaterally into the right vS1. Left: Epifluorescence image of a live coronal brain slice (300 μm) centered on vS2, from a P43 mouse injected in vS1 16 days prior with AAV2-ChR2-EYFP. (B) The recording schematic shows photostimulation of ChR2-expressing vS1 terminal arbors (green) and whole-cell recording from a non-expressing L2/3 RS neuron. Middle: Responses of a L2/3 RS cell in vS2 to intracellular current steps (scale bars 40 mV/ 200 ms). Right: vS1 excitatory synaptic current evoked in the same neuron (middle) by a pair of optical stimuli at 20 Hz (blue arrow, 0.5 ms) (average 13 sweeps). (C) Left: vS1 excitatory synaptic currents evoked in the same neuron (shown in B, middle) by a 20 Hz train of optical stimuli (average of 30 trials). Right: Average short-term dynamics of EPSCs evoked in L2/3 RS cells of vS2 during 20 Hz trains (n = 11 cells from 3 mice). Overall, long-range vS1-vS2 (sensory-sensory) CC responses facilitate similar to those observed in the vS1-vM1 (sensory-motor).

## DISCUSSION

Here we show that the synaptic properties of CC connections between vS1 and vM1 are dramatically different from other major excitatory connections of the neocortex. In contrast to local intracortical and core TC synapses, repetitive activation of many long-range CC synapses leads to short-term facilitation. Similar observations were made with a pathway linking cortical areas within the same sensory modality (vS1 to vS2), suggesting these properties may be a conserved feature of long-range corticocortical projections and not a unique feature of those between motor and sensory areas. A key difference between these pathways was that the synaptic dynamics were distinct for individual excitatory cells located across all cortical layers and specific subtypes of GABAergic neurons. We also found that the slow Ca^2+^ sensor Syt7 plays an important role in regulating facilitation at these synapses. A series of experiments further demonstrate that the overall excitation and spiking probability of L2/3 pyramidal neurons increases during repetitive stimulation of CC axons. This influence on pyramidal cell excitability depended on the distinct form of short-term plasticity at the CC synapse. Lastly, we identify several potential pitfalls of using optogenetic tools to study CC circuits that influence synaptic responses.

Short-term plasticity lasts from tens of milliseconds to minutes and is likely critical for information processing and cortical circuit function (Abbott et al. 1997; Tsodyks and Markram 1997; Abbott and Regehr 2004). Although the strength and dynamics of synapses targeting cortical neurons can vary widely (Stratford et al. 1996; Markram et al. 1998; Reyes et al. 1998; Gil et al. 1999; Beierlein et al. 2003; Brown and Hestrin 2009), these distinct properties may provide clues to their function and the type of information they convey. For example, core TC synapses are strong, reliable, and show robust short-term depression (Gil et al. 1999; Gabernet et al. 2005), perhaps reflecting the fidelity of the information they carry into the neocortex. On the other hand, the excitatory synapses between neighboring pyramidal cells within the cortex are typically weaker, less reliable, and show modest depression (Gil et al. 1999; Feldmeyer *et al*. 2002; Feldmeyer et al. 2006) (however see (Reyes and Sakmann 1999; Jouhanneau et al. 2015)). These synaptic features are likely important for how intracortical recurrent circuits distribute and combine information (Douglas et al. 1995; Lien and Scanziani 2013; Cossell et al. 2015; Cohen-Kashi Malina et al. 2016).

In contrast to TC and local CC excitatory synapses, most long-range CC connections displayed robust synaptic facilitation during repeated optical stimulation. This form of short-term plasticity is observed throughout the nervous system at synapses with a low probability of release (Jackman and Regehr 2017). However, this facilitation should not be confused with kinetic sluggishness since CC connections showed a strong ionotropic glutamate component in their response, suggesting an ability to carry rapid signals over sustained periods. The capacity to temporarily enhance their strength with activity suggests that CC synapses could efficiently convey information about ongoing local network activity to other cortical areas (Zagha et al. 2013). Short-term facilitation may also provide greater flexibility and control over target areas since their synaptic strength is activity-dependent, producing graded changes in excitation depending on the level of presynaptic spiking (Figure S3).

In vM1, vS1 responses were several times stronger and showed greater short-term facilitation in L2/3 than in other layers. The signals vM1 receives from vS1 are distinct from those conveyed to other cortical areas and likely encode object detection and location (Chen et al. 2013; Yamashita et al. 2013). L2/3 cells also supply a large fraction of the local excitatory drive to infragranular neurons that control motor output (Kaneko et al. 1994; Weiler et al. 2008; Hooks et al. 2011). Thus, these cells appear well-positioned to directly link sensory input and control of movement (Weiler et al. 2008; Mao et al. 2011; Sreenivasan et al. 2016). The facilitation patterns in the vS1 pathway suggest that L2/3 vM1 cells may be most excited during repeated spiking in vS1, such as those reported during repetitive sensory sampling or other active vibrissa-based behavioral tasks (Krupa et al. 2004; O’Connor et al. 2010; Vijayan et al. 2010).

We found that vM1 projections also engaged excitatory vS1 neurons with mostly facilitating synapses. The facilitation was particularly robust and sustained in L2/3 and L6, layers essential for associative interactions and modulating thalamocortical activities, respectively (Olsen et al. 2012; Lee et al. 2013; Crandall et al. 2015).

Although the two layers had similar facilitation, L6 responses were 3-4 times stronger, consistent with previous reports of a prominent input to this layer (Lee et al. 2008; Kinnischtzke et al. 2016; Zolnik *et al*. 2020). In addition, several studies have shown that vM1 activity correlates with various aspects of whisking (Carvell et al. 1996; Hill et al. 2011; Friedman et al. 2012; Castro-Alamancos 2013), and work by Petreanu et al. (2012) has demonstrated that vS1-targeting vM1 projection neurons increase their activity during vibrissa-dependent tasks, with some showing persistent-like activity.

Thus, the distinctive short-term dynamics of vM1 synapses may be a specialization tailored to inform vS1 neurons of behaviorally relevant features related to self-generated vibrissa movements over time.

In L5 of vS1, vM1 responses were relatively weak and showed different forms of synaptic plasticity, with approximately half of the inputs displaying depression. The disparate dynamics of these inputs imply at least two types of excitatory connections within L5, whose dynamic properties may be determined by the pre or postsynaptic cell type. The relatively weak responses could be due, in part, to the fact that many vM1 inputs target the apical tuft dendrites of L5 pyramidal neurons (Cauller et al. 1998; Petreanu et al. 2009), which make currents recorded at the soma prone to space-clamp errors and cable attenuation (Williams and Mitchell 2008). This issue suggests that distal vM1 inputs are underrepresented in the current study. Nonetheless, motor-related signals are very effective in modulating the activity of these neurons because they have powerful dendritic nonlinearities that help them actively integrate basal and apical tuft inputs when strong temporal correlations exist during active sensation and sensory perception (Xu et al. 2012; Larkum 2013; Manita et al. 2015; Takahashi et al. 2020).

CC synapse strength and dynamics also depend on the postsynaptic L2/3 interneuron subtype and pathway. For example, both vM1 and vS1 PV interneurons had the largest evoked currents of all cell types and displayed modest facilitation. In contrast, responses in vM1 and vS1 SOM cells were much weaker than neighboring RS cells but showed robust short-term facilitation. A different group of inhibitory interneurons, the VIP cells, had pathway-dependent strengths that depressed late in trains. VIP cells in vM1 had significantly weaker vS1-evoked responses than neighboring RS cells, whereas VIP interneurons in vS1 received similar amplitude vM1-evoked EPSCs. Although the latter finding is inconsistent with previous reports of vM1 synaptic currents being strongest in VIP cells (Lee et al. 2013; Naskar et al. 2021), we did find that these interneurons were very responsive to vM1 input, routinely firing action potentials (Figure S6). Thus, our data suggest that the greater propensity of VIP cells to spike in response to vM1 stimulation is due to differences in intrinsic membrane properties rather than synaptic strength (Figure S6). This mechanism is consistent with the idea that VIP interneurons are responsible for vM1-driven disinhibition of pyramidal neurons in vS1 (Lee et al. 2013).

ChR2 is widely used to stimulate axonal pathways to study neural circuits. However, we have identified several pitfalls associated with using optogenetic tools to study long-range CC connectivity. First, we found that some AAV serotypes used to express ChR2 can alter the short-term plasticity of CC synapses. Specifically, AAV1 and AAV5 impaired the magnitude of facilitation in a synapse-dependent manner, similar to observations in other systems (Jackman et al. 2014). Second, we observed AAV1, AAV5, and AAV9 transduce CC axon terminals more efficiently than AAV2, producing sparse but robust retrograde expression of ChR2 in cortical neurons that could potentially cause complications when interrogating these circuits. Third, we found that using the ChR2-mCherry fusion protein caused significant deficits in optically evoked synaptic plasticity, which may have been due to the aggregation of ChR2-mCherry in expressing neurons (Figure S7B), a reported problem when mCherry is expressed within some fusions (Asrican et al. 2013). Altogether, these AAV-induced changes to synaptic plasticity and the influence of mCherry tags on optogenetic probe expression and function could explain some of the discrepancies between the current study and previous work reporting depression (Lee et al. 2013; Kinnischtzke et al. 2014; Petrof et al. 2015; Zolnik et al. 2020; Naskar et al. 2021). However, in our hands, neither AAV serotype nor ChR2-mCherry alone caused short-term depression at CC synapses.

In addition to the optogenetic tool used, our data reveal that stimulation strategy and experimental conditions influence CC responses. Specifically, we show that the power density of over-terminal stimulation, light duration, and raising the concentration of extracellular Ca^2+^ decreased the magnitude of facilitation, likely because ChR2-induced presynaptic terminal depolarization and increased Ca^2+^ influx result in a greater release probability (Zucker 1999; Zhang and Oertner 2007; Jackman et al. 2014).

Although the power density used in previous studies is difficult to determine, the longer pulse durations (3-10 ms) and 2.0 mM extracellular Ca^2+^ concentration certainly contributed to the previously reported depressing CC responses (Lee et al. 2013; Kinnischtzke et al. 2014; Petrof et al. 2015; Zolnik et al. 2020; Naskar et al. 2021).

Here, we used a low power density, large light stimulation field to minimize complications that may arise due to direct stimulation of presynaptic terminals (Zhang and Oertner 2007; Jackman et al. 2014). Although this approach excites terminals, it probably stimulated CC axons away from their terminations as well. Axon stimulation is a more reliable way of activating presynaptic inputs in other pathways (Jackman et al. 2014) and is likely why robust facilitation was observed in this study. Consistent with this hypothesis, we found good agreement in the short-term plasticity evoked by our approach and over-axon stimulation. Although it is preferable to use over-axon stimulation, we found evoked synaptic responses using this method were considerably weak (<100 pA), probably due to the spatially diffuse nature of CC projections (Veinante and Deschenes 2003). The overall weaker evoked responses could complicate the study of synaptically driven circuit activity (i.e., disynaptic feedforward inhibition). Although the present approach appears to offer a reasonable strategy for studying the contributions of short-term plasticity to CC circuits, desensitization of ChR2 in presynaptic axons/terminals is still likely impacting some axons’ ability to follow during stimulation (Figure S2). Thus, CC synapses are probably more capable of maintaining sustained transmission than indicated by the data shown here.

One approach to overcome this problem and more reliably recruit axons during repetitive stimulation could be to employ opsins with kinetics faster than ChR2, such as Chronos (Klapoetke et al. 2014; Hass and Glickfeld 2016). However, not all faster opsins may be suitable. For example, Naskar et al. (2021) recently used ChETA, an engineered variant of ChR2 with fast kinetics (Gunaydin et al. 2010), to stimulate vM1-vS1 projections, and reported significant short-term depression. Thus, assessing the fidelity of optogenetic activation in a given pathway is still critical when using any opsin.

Our data begin to unravel how synaptic dynamics may contribute to long-range CC influences. Obviously, synaptic dynamics are not the only factor determining the net CC influence on target areas. Previous work has emphasized crucial roles for topographic alignment, interareal synchrony, network state, synapse distribution, disinhibition, as well as active membrane properties of pyramidal cells (Hoffer et al. 2003; Petreanu et al. 2009; Xu et al. 2012; Larkum 2013; Lee et al. 2013; Zagha et al. 2013; Bastos et al. 2014; Manita et al. 2015). Furthermore, trans-thalamic pathways have been proposed to contribute to CC communication (Sherman and Guillery 2011). The current work complements these elegant studies, and, together, they indicate the dynamic state of CC synapses has an essential role in ongoing cortical functioning.

### SUPPLEMENTAL MATERIAL

Supplemental material is available online.

## FUNDING

This work was supported by the National Institutes of Health (NIH) K99-NS096108 (to S.R.C), R00-NS096108 (to S.R.C), and R00-NS096108-S1 (to S.R.C and L.E.M).

### NOTES

We thank Barry Connors (Brown University), Scott Cruikshank (The University of Alabama at Birmingham), and Suryadeep Dash (Crandall Lab) for helpful discussions and comments on the manuscript. We thank Barry Connors for generously providing the *Ntsr1-Cre*, *Rbp4-Cre*, and *5HT3a-EGFP* mice. We thank Charles Lee Cox (Michigan State University) for imaging support. We also thank the Michigan State University Center for Advanced Microscopy for confocal imaging support. *Conflict of Interest*: None declared.

### AUTHOR CONTRIBUTIONS

S.R.C designed the experiments and supervised the study; L.E.M., K.E.B., H.H.K, and S.R.C performed the electrophysiological experiments; D.M.A performed the histological experiments; L.E.M., K.E.B., H.H.K, and S.R.C analyzed the data; S.R.C. prepared and wrote the manuscript. L.E.M., K.E.B., H.H.K, and D.M.A reviewed and edited the manuscript. All authors approved the final manuscript.

## MATERIALS AND METHODS

### Animals

All procedures were carried out in accordance with the National Institutes of Health (NIH) Guidelines for the Care and Use of Laboratory Animals and approved by the Michigan State University Institutional Animal Care and Use Committee (IACUC). We used the following mouse lines in this study: Crl: CD1 (ICR) (Charles River: 022), Ai14 (Jackson Labs: 007908) (Madisen et al. 2010), *PV-Cre* (Jackson Labs: 008069) (Hippenmeyer et al. 2005), *SOM-IRES-Cre* (Jackson Labs: 013044) (Taniguchi et al. 2011), *VIP-IRES-Cre* (Jackson Labs: 010908) (Taniguchi et al. 2011), *Scnn1a-Tg3-Cre* (Jackson Labs: 009613) (Madisen et al. 2010), *Syt7* knockout (Jackson Labs: 004950) (Chakrabarti et al. 2003), *Ntsr1-Cre* (MMRRC: 017266-UCD) (Gong et al. 2007), *Rbp4-Cre* (MMRRC: 031125-UCD) (Gong et al. 2007), and *5HT3a-EGFP* (MMRRC: 000273-UNC). The *Ntsr1-Cre*, *Rbp4-Cre*, *Scnn1a-Tg3-Cre*, *Syt7* knockout, and *5HT3a-EGFP* mouse lines had ICR genetic backgrounds. All mice, except for ICR, *Syt7* knockout, and *5HT3a-EGFP* mice, were bred by crossing homozygous or heterozygous Cre mice with homozygous Ai14 reporter mice, resulting in experimental mice that were heterozygous for the indicated genes. Animals were group-housed with same-sex littermates in a dedicated animal care facility maintained on a 12:12 hour light-dark cycle. Food and water were available *ad libitum*. We used both male and female mice in this study.

### Stereotactic Virus Injections

For all functional experiments characterizing the properties of CC synapses, we used an adeno-associated virus (AAV2) that encoded genes for hChR2 (H134R)-EYFP fusion proteins (rAAV2/hSyn-hChR2[H134R]-eYFP-WPREpA, AV4384, University of North Carolina Viral Vector Core). For the AAV serotype experiments, we also used the following AAV vectors obtained from Addgene: pAAV1-hSyn-hChR2(H134R)-EYFP, pAAV5-hSyn-hChR2(H134R)-EYFP, and pAAV9-hSyn-hChR2(H134R)-EYFP (Addgene plasmid # 26973). For the fusion protein experiment, we used AAV2/hSyn-hChR2(H134R)-mCherry (AV4385, University of North Carolina Viral Vector Core).

Retrograde-Cre experiments were carried out similarly, with injections of AAVretro-Ef1a-mCherry-IRES-Cre (Addgene plasmid # 55632) into vS1 followed by an AAV injection in vM1 that drove Cre-dependent expression of ChR2 (pAAV1-hSyn-hChR2(H134R)-EYFP, Addgene plasmid # 26973). We performed dual-optogenetic experiments with injections of rAAV2/hSyn-hChR2[H134R]-eYFP-WPREpA (AV4384, University of North Carolina Viral Vector Core) into vS1 followed by injecting AAV2/hsyn-Flex-ChrimsonR-tdT (AV6555, University of North Carolina Viral Vector Core) into vM1 1-week after of a *PV-Cre* mouse. We achieved selective optical control of local excitatory neurons and axons/terminals by injecting a Cre-dependent AAV2 in Cre-driver mice (*Scnn1a-Tg3-Cre* and *Rbp4-Cre*) (rAAV2/EF1a-DIO-hChR2(H134R)-eYFP, AV4378, University of North Carolina Viral Vector Core).

For all surgeries, the virus was injected unilaterally into vS1 or vM1 of mice *in vivo*, as previously described (Crandall *et al*. 2015; Crandall *et al*. 2017). Injections were normally performed on mice ∼3 weeks old (Mean injection age: 22.2 ± 0.7 days, range 17-42 days). Briefly, mice were anesthetized with a Ketamine-Dexdomitor mixture diluted in sterile saline (KetaVed, 70 mg/kg; Dexdomitor, 0.25 mg/kg; intraperitoneally). Once deeply anesthetized, mice were placed into a digital stereotaxic frame with an integrated warming base that maintained core body temperature (Stoelting). A thin layer of ophthalmic ointment was applied to the eyes to prevent drying (Patterson Veterinary Artificial Tears). Next, an incision was made over the skull by scalpel or fine surgical scissors, the scalp and periosteum overlying the skull deflected, and a small craniotomy made over the target site. A small volume of the virus was then pressure-ejected via a glass micropipette attached to a Picospritzer pressure system (0.1-0.2 μl per injection site over 5-10 min; titer = 3.1-3.5 x 10^12^ viral genomes/ml). In some experiments, we used a virus-retrobead or saline-retrobead mixture (red RetroBeads, Lumafluor, Cat# R180). When comparing different AAV serotypes, we kept titer levels consistent by diluting each viral preparation to the same titer (3.1 x 10^12^ viral genomes/ml) just before intracranial injection. Following injection, the pipette was held in place for an additional 5-10 min before being slowly advanced or withdrawn from the brain. After surgery, the scalp was closed with a surgical adhesive (GLUture), and animals were given Antisedan (2.5 mg/kg) to reverse the effects of Dexdomitor. Mice were allowed to recover on a heating pad for 1 hr before returning to their home cage. Most experiments were performed ∼3 weeks after the virus injections to allow sufficient expression (Mean expression time: 20.4 ± 0.3, range 11-25 days). Coordinates from bregma for vS1 were 3.4 mm lateral, 0.8 mm posterior, 0.40, and 1.0 mm depth. Coordinates for vM1 were 1.25 mm lateral, 0.9 and 1.3 mm anterior, 0.40, and 1.0 mm depth. Coordinates for VPm were 1.8 mm lateral, 0.75 mm posterior, 3.05 mm depth.

### *In Vitro* Slice Preparation

After allowing ∼3 weeks for ChR2-EYFP expression (Mean experimental age: 44.0 ± 3.3 days, Median age: 41 days, range 30-259 days), acute coronal brain slices (300 μm thick) containing vM1 and vS1 were prepared for *in vitro* recording, as described previously (Crandall et al. 2010; Crandall *et al*. 2017). In four mice, acute thalamocortical brain slices (300 μm thick) containing VPm and vS1 were prepared, as described previously (Agmon and Connors 1991; Crandall et al. 2017). Briefly, animals were anesthetized with isoflurane before being decapitated. Brains were removed and placed in a cold (∼4° C) oxygenated (95% O_2_, 5% CO_2_) slicing solution containing (in mM) 3 KCl, 1.25 NaH_2_PO_4_, 10 MgSO_4_, 0.5 CaCl_2_, 26 NaHCO_3_, 10 glucose, and 234 sucrose. Brain slices were cut using a vibrating tissue slicer (Leica, VT1200S) and then transferred to a holding chamber with warm (32° C) oxygenated artificial cerebrospinal fluid (ACSF) containing (in mM): 126 NaCl, 3 KCl, 1.25 NaH_2_PO_4_, 2 MgSO_4_, 2 CaCl_2_, 26 NaHCO_3_, and 10 glucose. Slices were maintained at 32° C for 20 min and then at room temperature for a minimum of 40 min before recording. Slices containing the injection site were always collected for imaging to confirm injection site accuracy and assess tissue health. We only considered mice in which there were no signs of tissue damage or off-target injections.

### *In Vitro* Electrophysiological Recordings and Data Acquisition

Individual brain slices (300 μm) were transferred to a submersion-type recording chamber and bathed continually (2.5 - 3.0 ml/min) with warm (32 ± 1°C) oxygenated ACSF containing (in mM): 126 NaCl, 3 KCl, 1.25 NaH_2_PO_4_, 1 MgSO_4_, 1.2 CaCl_2_, 26 NaHCO_3_, and 10 glucose. Neurons were visualized using infrared differential interference contrast optics (IR-DIC) with a Zeiss Axio Examiner.A1 microscope equipped with a 40x water immersion objective (Zeiss, W Plan-Apo 40x/1.0NA) and video camera (Olympus, XM10-IR). Whole-cell recordings were obtained using patch pipettes with tip resistances of 4-6 MΩ when filled with a potassium-based internal solution containing (in mM): 130 K-gluconate, 4 KCl, 2 NaCl, 10 HEPES, 0.2 EGTA, 4 ATP-Mg, 0.3 GTP-Tris, and 14 phosphocreatine-K (pH 7.25, 290 mOsm). Voltages were corrected for a −14 mV liquid junction potential. The average reversal potential for GABA_A_ receptor-mediated inhibitory responses was −91 mV when measured in excitatory cortical cells (n = 3 cells). The calculated reversal potential was −95 mV. Neurobiotin (0.25%, w/v; Vector Laboratories) was added to the internal solution to inject into neurons during whole-cell recordings for subsequent identification in a subset of experiments (Figure S3).

Electrophysiological data were acquired and digitized at 20 kHz using Molecular Devices hardware and software (MultiClamp 700B amplifier, Digidata 1550B4, pClamp 11). Signals were low-pass filtered at 10 kHz (current-clamp) or 3 kHz (voltage-clamp) prior to digitizing. During recordings, the pipette capacitances were neutralized, and series resistances (typically 10-25 MΩ) were compensated online (100% for current-clamp and 70-80% for voltage-clamp). Series resistances were continually monitored and adjusted during experiments to ensure sufficient compensation. The local field potential (LFP) was monitored with a patch-style pipette (∼0.6 MΩ) filled with 3 M NaCl, and signals were band-pass filtered between 0.1 Hz – 4 kHz. The LFP pipette was placed in L1 for recordings in vS1 and L2/3 for recordings in vM1. Cell attached recordings were obtained using patch-style pipettes filled with a potassium-based internal solution (see above), and signals were high pass filtered (100 Hz). All pharmacological agents were bath-applied at least 10 min before subsequent experimental tests. Pharmacological agents included Tetrodotoxin citrate (TTX, Tocris, Cat# 1069), 4-Aminopyridine (4-AP, Sigma, Cat# A78403), DL-AP5 (Sigma, Cat# A5282), DNQX (Sigma, Cat# D0540), Picrotoxin (Sigma, Cat# P1675), and CGP 55845 hydrochloride (Tocris, Cat# 1248).

### Photostimulation

ChR2 was optically excited using a high-power white light-emitting diode (LED) (Mightex, LCS-5500-03-22) driven by a LED controller (Mightex, BLS-1000-2). For the two-color photostimulation experiments (Figure S4), ChR2 and Chrimson were excited using a high-power blue (455 nm; LCS-0455-03-22) and red (625 nm; LCS-0625-03-22) LED, respectively, combined by a multi-wavelength beam combiner (Mightex, LCS-BC25-0495). LED on/off times were fast (< 50 μs) and of constant amplitude and duration when verified with a fast photodiode (Thorlabs, DET36A). The light was collimated and reflected through a single-edge dichroic beam-splitter (Semrock, FF660-FDi02) and a high-magnification water immersion objective (Zeiss, W Plan-Apo 40x/1.0 NA), resulting in an estimated spot diameter of ∼1500 µm and maximum white LED power at the focal plane of ∼32 mW (∼18 mW/mm^2^). Stimuli were delivered as 0.5 ms flashes and were directed at ChR2-expressing axons/terminals by centering the light over the recorded cell. The LED intensity was typically adjusted to evoke a 200 pA EPSC in the recorded neuron when held in voltage-clamp at −94 mV, near the reversal potential for inhibition (see Table S1 for the LED intensities needed for each cell type). This intensity was always subthreshold for the recorded cell (∼3 mV; Table S1). For the over terminal stimulation experiments (Figure 6), ChR2 was excited using a high-power white LED (Mightex, BLS-GCS-6500-65-A0510) attached to a Digital Mirror Device (DMD) based pattern illuminator (Mightex, Polygon 400). The pattern illuminator was used to briefly illuminate a small spot (0.5-ms flashes; 150 μm diameter) centered over the soma (Max LED power across all mirrors ∼44.3; Max LED power when illuminating a 150 μm diameter spot ∼4.9 mW or 276 mW/mm^2^). For these experiments, the LED light was reflected through the same water immersion objective as above.

### Live Slice Imaging

Before recording, all live brain slices (300 μm) were imaged using a Zeiss Axio Examiner.A1 microscope equipped with a 2.5x objective (Zeiss, EC Plan-Neofluar) and Olympus XM10IR camera and cellSens software. Brightfield and fluorescence images were obtained to confirm the accuracy of the injection, tissue health at the injection site, and the overall expression in axons/terminals in the target region. During imaging, live slices were kept in a submersion recording chamber and continually bathed with oxygenated ACSF (at room temperature).

### Immunohistochemistry and Tissue Processing for Neurobiotin

All tissue for immunohistochemistry was prepared from acute coronal brain slices, as described previously (Crandall et al. 2017) (Mean age of genotype pairs: 97.8 ± 9.3 days, range 73-116 days). Briefly, acute brain slices (300 μm) containing vS1 or vM1 were fixed by immersion overnight at 4°C in a vial containing 4% paraformaldehyde in 0.1 M phosphate buffer (PB) solution. Slices were then transferred to a 30% sucrose in M PB solution until re-sectioned (4°C; 2-3 days). Tissue was re-sectioned at 60 μm using a freezing microtome (Leica, SM2010 R). Next, sections were immunostained for Syt7, as described previously (Jackman *et al*. 2016; Turecek et al. 2017). Briefly, sections were washed 2 times in 0.1 M phosphate buffer followed by 3 times in 0.1 M phosphate buffer with 0.9% NaCl, pH 7.4 (PBS) (5 min per wash). After washing, sections were pre-incubated for 1 hour at room temperature with a blocking solution (0.1% Tween, 0.25% Triton X-100, 10% normal goat serum in PBS), then incubated with primary antibody for 24 hours with rotation at 4°C. Following primary incubation, sections were washed 5 times in PBS (5 min per wash), pre-incubated for 45 min in blocking solution (same as above), incubated with secondary antibody for 2 hours at room temperature, then washed 3 times in PBS (10 min per wash) and 2 times in PB (5 min per wash). Sections were mounted and cover-slipped using Vectashield with DAPI (Vector Laboratories H-1500). The primary antibody was rabbit polyclonal anti-synaptotagmin 7 (1:200, Synaptic Systems, Cat# 105-173), and the secondary antibody goat anti-rabbit IgG (H+L) cross-adsorbed secondary antibody Alexa Fluor 568 (1:500, Thermo Fisher Scientific, Cat# A-11011). Antibodies were stored and prepared as recommended by the manufacturer and did not undergo repeated freeze/thaw cycles. For experiments comparing genotypes (WT and Syt7 KO), all tissue was prepared, stained, and processed in parallel. For each staining run, no primary and no secondary controls were conducted. Images were taken using a Zeiss Axio Imager.D2 fluorescence microscope equipped with appropriate filter sets and a high-resolution monochrome digital camera (Zeiss, AxioCam) and Zen software. For each genotype pair, images were collected from identical anatomical locations, using the same microscope settings, and processed identically in Fiji: ImageJ.

Slices containing cells injected with neurobiotin were transferred to 4% paraformaldehyde in 0.1 M PB solution overnight at 4°C (18 – 24 hr). The next day, slices were changed to 30% sucrose in 0.1 M PB until re-sectioned (4°C; 2 – 3 days). Individual brain slices were re-sectioned at 100-150 µm using a freezing microtome (Leica SM2010 R). Floating sections were washed twice in 0.1 M PB followed by three washes in 0.1 M PBS, pH 7.4 (5 min per wash). After washing, sections were incubated for 1 hr at room temperature in a blocking solution containing 0.1% Tween, 0.25% Triton X-100, and 10% normal goat serum in PBS. Sections were then incubated using Streptavidin/Biotin Blocking Kit (Vector Labs, SP-2002), 30 minutes in streptavidin solution, followed by 30 minutes in biotin solution with a brief rinse of PBS after each.

Sections were then incubated with Alexa Fluor 568-conjugated streptavidin (Thermo-Fisher Scientific, S11226, 1:5000, 2μg/μl) solution in blocking solution for 3 hrs with rotation at room temperature. Following incubation, sections were washed 3 times in PBS, then 2 times in 0.1 M PB solution (10 min per wash), mounted, and cover-slipped using Vectashield Vibrance with DAPI (Vector Laboratories, H1800). Confocal image stacks of labeled neurons were taken using an Olympus FluoView 1000 filter-based Laser Scanning Confocal Microscope with an Olympus 40x Oil UPLFLN O (NA 1.3) objective and updated Version 4.2 software (laser excitation 543nm).

### Electrophysiological Data Analysis

Analysis of electrophysiological data was performed in Molecular Devices Clampfit 11 and Microsoft Excel. Synaptic responses to optical stimulation were measured from postsynaptic neurons recorded in whole-cell voltage-clamp. The amplitude of an evoked EPSC was measured relative to a baseline before the stimulus (0.5-10 ms depending on the frequency of the stimulation). The EPSC peak was measured over the 10 ms immediately after stimulus onset or over the 4.5 ms immediately after stimulus onset for all layer comparisons. Reported values were based on average responses to 3–20 stimuli (typically 10). Values for the fiber volley and opsin potential were based on average responses to ∼200 stimuli, and their peaks were measured over the 10 ms immediately after stimulus onset. The field postsynaptic potential (fPSP) slope was measured over a 0.5 ms region immediately after the peak negativity. Intrinsic physiological properties were measured using previously described methods (Crandall et al., 2017). Regular-spiking (RS) neurons were identified using established physiological criteria, including their spike-frequency adaptation, growing afterhyperpolarization (AHP) during spike trains, and relatively long spike half-widths (McCormick et al. 1985). In a subset of experiments, we confirmed that RS cells were excitatory (pyramidal) neurons (Figure S3F-H).

### Layer Comparisons Using Cre-Driver Mouse Lines

Previous work has shown that the Cre-driver mouse lines used in this study are selective for excitatory cells of distinct layers (L4: *Scnn1a-Tg3-Cre*; L5: *Rbp4-Cre*; L6: *Ntsr1-Cre*) (Olsen *et al*. 2012; Kim et al. 2014; Adesnik 2018). Live slices prepared from Cre mice crossed with a tdTomato (tdT) reporter (Ai14) confirmed that Cre-expressing cells were mostly confined to their respective layers and that the subpial distances for the layers were in agreement with previous reports (Table S3). Layers were identified by drawing horizontal lines where the density of tdTomato-expressing cells decreased (Frandolig et al. 2019). Depth measurements for cells are reported as absolute distance or normalized distance from the top of their respective layer. Whole-cell recordings further confirmed that all tdTomato positive cells had intrinsic physiological properties consistent with excitatory cortical cells (McCormick et al. 1985; Crandall et al. 2017). No Cre expression was observed in vM1 of the *Scnn1-Cre* mouse, consistent with the idea that the agranular cortex lacks spiny stellate cells (Harris and Shepherd 2015). A Cre-driver mouse line was not used for L2/3 recordings. However, all L2/3 cells were confirmed posthoc to be located within the superficial layers of the cortex (average recorded depth from pia: vS1 L2/3 cells, 190.5 ± 3.9 μm, range: 120-310 μm; vM1 L2/3 cells: 238.0 ± 5.4 μm, range: 135-361 μm). To control for variability in ChR2 expression in different slices and mice, we sequentially recorded from an identified L2/3 RS cell and an excitatory neuron (tdTomato-positive and tdTomato-negative) in a separate layer of the same column. For synaptic strength comparisons, brief flashes of the same intensity were delivered over the recorded soma.

### Experimental Design and Statistical Analysis

No formal method for randomization was used in this study. Experiments and data analyses were performed blind to the conditions of the Syt7 experiments. For all other studies, the experimenters were not blind to experimental groups. No statistical methods were used to predetermine sample size, but our sample sizes are similar to those reported in previous studies (Crandall et al. 2015; Crandall et al. 2017). Statistical comparisons were performed in OriginPro 2019. The Shapiro-Wilk test was first applied to determine whether the data had been drawn from a normally distributed population, in which case parametric tests were used. If the assumption of normality was not valid, non-parametric tests were used. Significance was assessed using the appropriate parametric (Paired t-test or Two-sample t-test) or non-parametric test (Wilcoxon paired signed-rank test or Mann-Whitney U test) as indicated in the Results. A two-way analysis of variance (ANOVA) was used to compare short-term synaptic dynamics across 20 Hz trains. A one-way ANOVA was used for multiple comparisons. All tests were two-tailed. Data are represented as mean ± SEM, and statistical significance was defined as p < 0.05.

## SUPPLEMENTAL FIGURE LEGENDS

**Figure S1.**
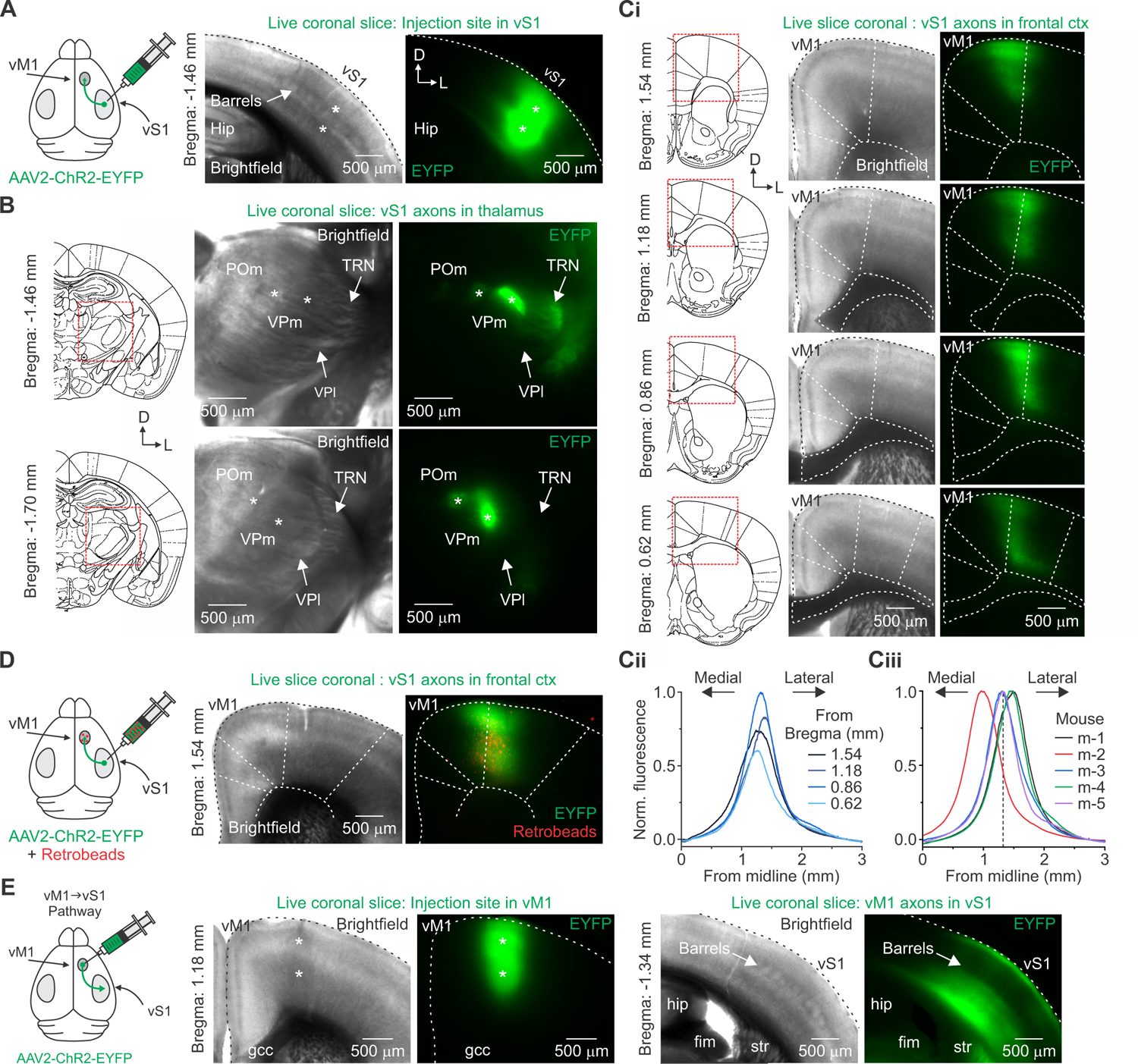
Mapping the region of the frontal cortex that mediates long-range reciprocal connections between vS1 and vM1. Related to **Figure 1**. (A) Left: AAV2-ChR2-EYFP was injected into vS1 of mice *in vivo*, and then live 300 μm-thick serial coronal sections were collected 20-22 days after for imaging. Right: Brightfield and epifluorescence images of a live coronal slice through vS1 of a P40 mouse. The presence of L4 barrels identified vS1. Asterisks indicate the viral injection sites. (B) Left and middle: Reference atlas drawings and brightfield images of consecutive coronal slices through somatosensory thalamus (same mouse as A). Right: Epifluorescence images show EYFP-expressing axons/terminals in the somatosensory thalamus, including the thalamic reticular nucleus (TRN), ventral posterior medial nucleus (VPm), and posterior medial nucleus (POm). No expression was observed in the ventral posterior lateral nucleus (VPl). (C) Top (Ci): Reference atlas drawings and serial slices through frontal cortex (same mouse as A). EYFP-expressing vS1 projections formed a narrow band just lateral to midline that was strongest between 0.5 and 2.0 mm anterior to bregma when compared to a reference atlas. (Cii), Fluorescence profile plot for each image shown in Ci, normalized to the strongest expressing slice. The anterior-posterior coordinate for vM1 was based on the approximate location of the strongest expressing slice. (Ciii), Normalized fluorescence profile plot of the strongest expressing slice for each mouse. The distance of the peak from the midline was used to calculate the medial-lateral coordinate for vM1. (Average vM1 coordinates: anterior-posterior location from bregma, 1.12 ± 0.13 mm; medial-lateral location from midline, 1.27 ± 0.07 mm; half-width of the normalized fluorescence plot, 0.65 ± 0.02 mm; n = 5 mice). (D) vM1 and vS1 are reciprocally connected. Left: AAV2-ChR2-EYFP and retrobeads were co-injected into vS1. Right: Brightfield and epifluorescence image of a live coronal slice through vM1 of a P39 mouse (23 days of expression). Shown is the overlap of retrograde cells (red) and vS1 anterograde axons/terminals (green) in vM1 (results were repeated in n = 4 mice). (E) vM1 axons terminate diffusely in vS1. Left: AAV2.ChR2.EYFP was injected unilaterally into vM1 using the coordinates above. The virus serotype, volume, and the location of each injection (see Methods) were chosen to best match the pattern of vS1 terminal arbors and to minimize spread into neighboring areas (compare the vM1 images in E with C and D). Right: Brightfield and epifluorescence images through vM1 and vS1 of a P39 mouse injected in vM1 21 days prior with AAV2-ChR2-EYFP. The vS1 images are used in Figure 1. Values are represented as mean ± SEM.

**Figure S2.**
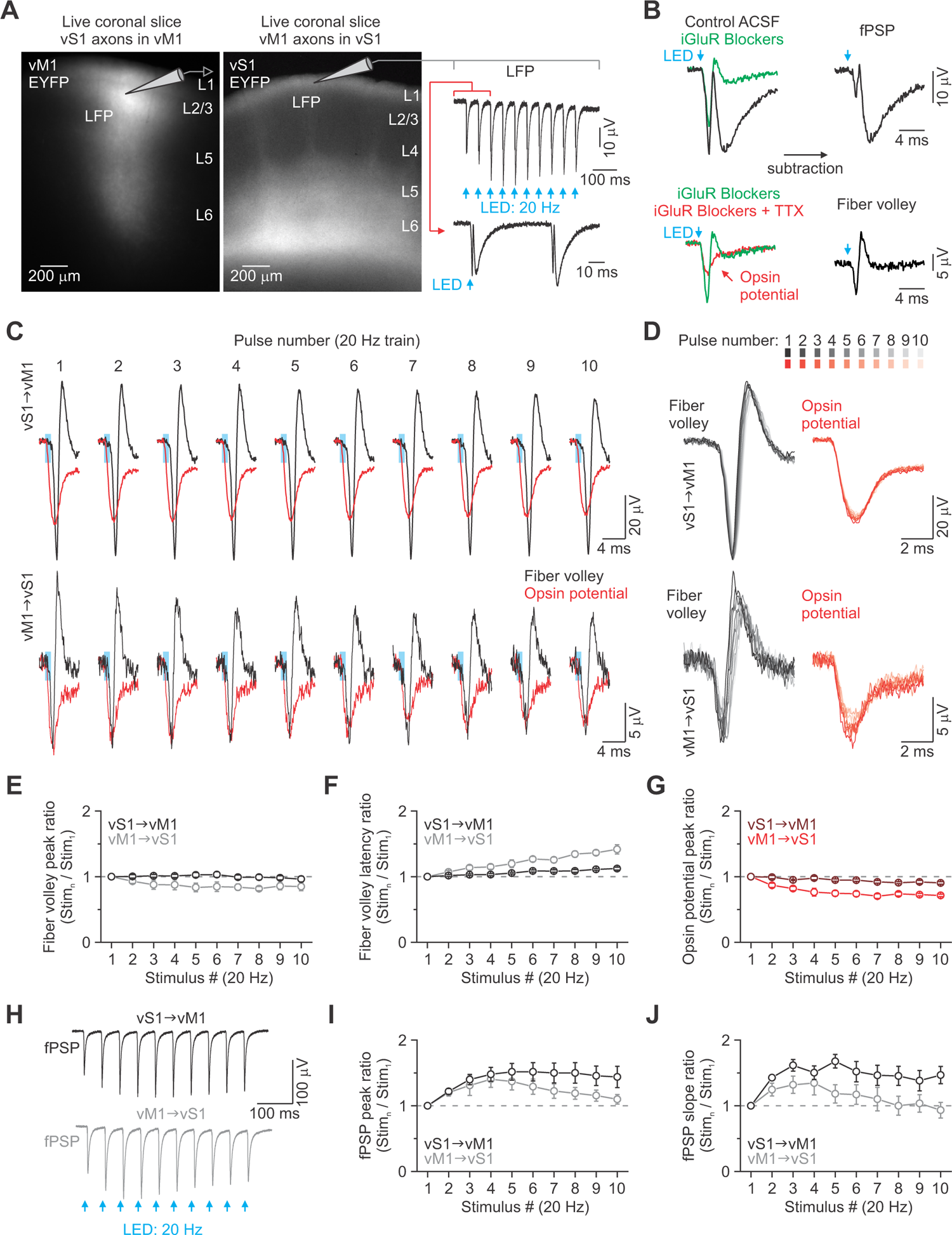
ChR2 can reliably drive action potentials in long-range CC axons. Related to Figure 1. (A) Left: Epifluorescence image of a live 300 μm coronal slice showing ChR2-EYFP expressing vS1 terminal arbors in vM1 of a P39 mouse (22 days after viral injection in vS1). Middle: Similar image showing ChR2-EYFP expressing vM1 terminal arbors in vS1 of a P41 mouse (21 days after viral injection in vM1). Right: To test the effectiveness of ChR2 in driving presynaptic action potentials (APs), we recorded the local field potential (LFP) in response to a 20 Hz train (10 pulses, 0.5 ms pulse duration, every 10 sec). LFP recordings and optical stimulation were targeted to L2/3 and L1 for vM1 and vS1, respectively. Light intensities were adjusted for each slice to the power needed to evoke a 200 pA EPSC in a neighboring L2/3 RS cell when held near the inhibitory reversal potential (average LED power: 1.7 mW or 1.0 mW/mm^2^ for vS1 axons in vM1, 12.4 mW or 7.0 mW/mm^2^ for vM1 axons in vS1). (B) Example traces showing how the fiber volley was pharmacologically isolated from the recorded LFP (Hass and Glickfeld 2016). Ionotropic glutamate receptors (iGluRs) were first blocked by adding selective antagonists (DNQX, 20 μM; APV, 50 μM) to the control ACSF. Subtracting the resulting trace from the control produced the field postsynaptic potential (fPSP). Sodium channels were subsequently blocked by adding tetrodotoxin (TTX, 1 μM). The resulting TTX-insensitive potential was referred to as the opsin potential (Hass and Glickfeld 2016). Subtracting the opsin potential from the potential recording in iGluR blockers resulted in the isolated fiber volley (bottom right). (C) Repetitive activation of CC axons with ChR2 is reliable. Fiber volleys (black) and opsin potentials (red) evoked by a 20 Hz train of optical stimuli (blue boxes: 0.5 ms) for both the vS1-vM1 (top) and vM1-vS1 (bottom) pathways (average of ∼200 trials each). (D) Overlay of the fiber volleys and opsin potentials from the recordings in (C). (E-G) Population data showing the normalized fiber volley amplitude (E), fiber volley latency (F), and opsin potential amplitude (G) during 20 Hz trains (n = 10 slices from 5 mice for the vS1-vM1 pathway; n = 7 slices from 4 mice for the vM1-vS1 pathway). Repetitive 20 Hz stimulation of ChR2-expressing vS1 axons/terminals resulted in a fiber volley amplitude decay of ∼0% after the second pulse and 4% on the tenth pulse. The same stimulation of ChR2-expressing vM1 axons/terminals resulted in a modest fiber volley amplitude decay of ∼7% after the second pulse and ∼15% on the tenth pulse (E). (H) Isolated fPSPs evoked in vM1 and vS1 by a 20 Hz train of optical stimuli (average of 200 trials each). (I-J) Population data showing the normalized fPSP amplitude and fPSP slope during 20 Hz trains (n = 10 slices from 5 mice for the vS1-vM1 pathway; n = 7 slices from 4 mice for the vM1-vS1 pathway). Repetitive 20 Hz stimulation resulted in modest facilitation in fPSP amplitude and slope for both pathways. Values are represented as mean ± SEM.

**Figure S3.**
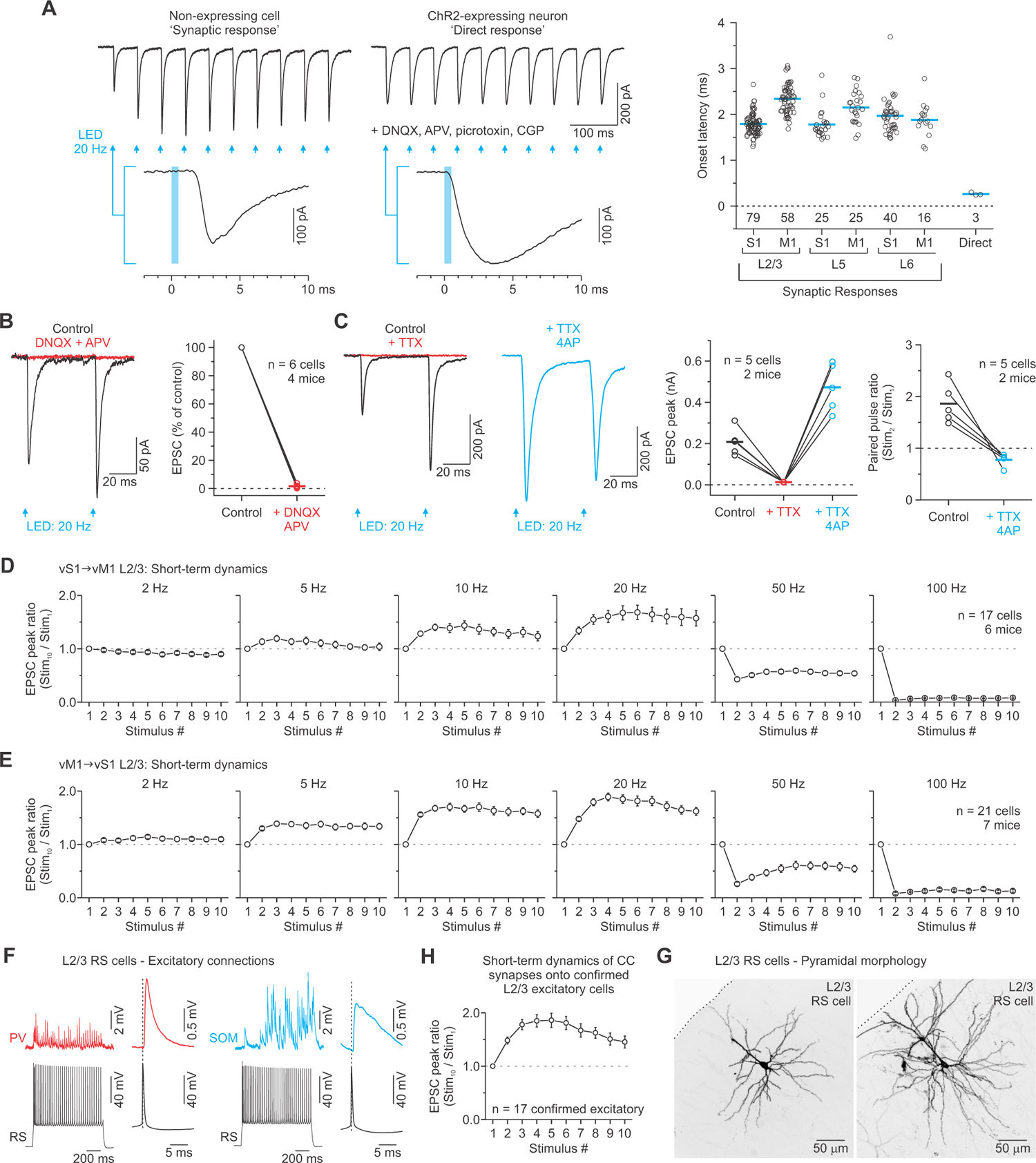
Quantification of onset latencies, glutamate sensitivity, action potential dependence, and frequency dependence of ChR2-evoked responses in L2/3 of the vS1-vM1 CC system. Related to Figure 1. (A) Optically evoked synaptic responses had delays consistent with the timing of neurotransmission at fast synapses. Left: Current responses of a non-expressing L2/3 RS cell and a ChR2-expressing L2/3 RS cell to brief wide-field LED illumination (10 pulses, 20 Hz, 0.5 ms pulse duration). An expanded trace of the first response shows a 1.8 ms response delay from the non-expressing cell (synaptic response) and a 0.24 ms response delay for the expressing cell (direct response) (average of 10 and 13 trials). During the recordings of ChR2-expressing cells, glutamate and GABA receptor antagonists were bath applied to block synaptic transmission (DNQX, 20 μM; APV, 50 μM; picrotoxin, 50 μM; CGP-55845, 2 μM). Right: Population plot of the onset latencies for the vS1 and vM1 synaptic responses and ChR2-expressing cortical neurons (direct). Synaptic responses are divided according to the cortical and laminar location of the recorded cell. Cell counts are located below the zero line. Each point represents the average onset latency for an individual neuron, measured with a LED intensity adjusted to obtain a 200 pA EPSC. Blue lines indicate average latencies for each population. Note that the onset latencies of ChR2-expressing neurons were significantly faster than synaptic responses, consistent with previous reports (Cruikshank et al. 2010; Crandall et al. 2017). The average onset latency for the entire population (excluding L4 in vS1) was 1.99 ± 0.02 ms (n = 239 cells). (B) Optically evoked CC synaptic responses require fast glutamatergic transmission. Left: In control ACSF, synaptic currents evoked in an RS cell by a pair of 0.5 ms optical stimuli at 20 Hz (black trace) (average of 10 trials). Bath application of the glutamate receptor antagonists APV (50 μM) and DNQX (20 μM) for 10 min blocked >98% of the postsynaptic current (red trace) (average of 10 trials). Right: Population effects of glutamate receptor antagonists on LED-evoked CC responses (n = 6 cells from 4 mice). (C) Optically evoked CC synaptic responses require presynaptic APs and are monosynaptic. Left: In control ACSF, synaptic currents were evoked in an RS cell by a pair of 0.5 ms optical stimuli at 20 Hz (black trace) (average of 14 trials). Bath application of the sodium channel blocker TTX (1 μM) for 10 min blocked 100% of the postsynaptic current (red trace) (average of 14 trials) (similar results seen in n = 5 cells from 2 mice). Middle: For the same cell, subsequent bath application of the potassium channel blocker 4-AP (1 mM) for 10 min rescued the EPSC (blue trace) (average of 14 trials) (Petreanu et al. 2009). The currents evoked in TTX+4AP were larger, slower, and the dynamics switched from facilitating to depressing. Right: Population effects of TTX and TTX+4AP on optically evoked CC response amplitudes (n = 5 cells from 2 mice). Population effect of TTX+4AP on the paired-pulse ratio of CC responses (n = 5 cells from 2 mice). The horizontal bars are averages for each group. (D-E) Facilitation is present in long-range CC synapses at multiple frequencies. Population plots of the average normalized peak synaptic responses evoked by 2-100 Hz optical trains (10 stimuli; 0.5 ms pulse duration). Results for both the vS1-vM1 (D) and vM1-vS1 (E) pathways are shown. Synaptic enhancement during trains was present over a range of frequencies. Responses switched to depression during 50 and 100 Hz trains, most likely due to the properties of ChR2 (Lin et al. 2009). (F) Confirmation that RS neurons are excitatory. EPSPs recorded from PV and SOM cells, evoked by stimulation of presynaptic L2/3 RS cells. Inset shows a spike-triggered average EPSP. (G) Summary data of CC-evoked EPSC dynamics from RS cells confirmed to be excitatory (n = 17 cells from 15 mice). (H) Confocal image (Z projection) of two RS cells filled with neurobiotin. Their pyramidal shape confirms they were excitatory neurons. Values are represented as mean ± SEM.

**Figure S4.**
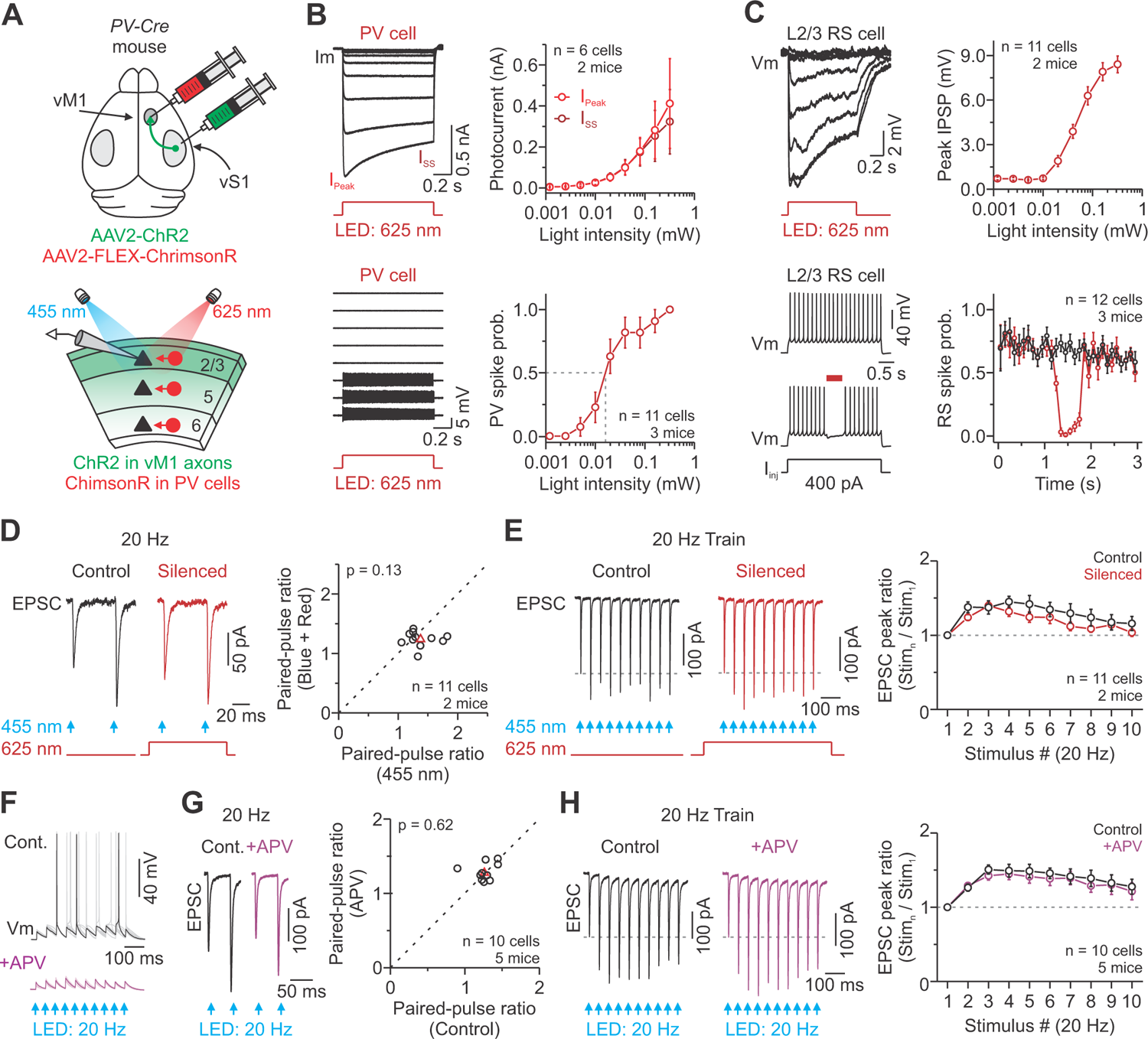
Optogenetically or pharmacologically silencing local excitatory connections reveals intracortical inputs do not confound long-range CC facilitation. Related to Figure 1. (A) Schematic illustration of cortical silencing experiments to determine if intracortical excitatory inputs confound long-range CC facilitation. Top: We injected AAV2-ChR2-EYFP and AAV2-Flex-ChrimsonR into the cortex of *PV-Cre* mice *in vivo,* and then live 300 μm-thick coronal brain sections were collected for *in vitro* recordings. Bottom: We optogenetically silenced local excitatory cells by activating ChrimsonR-expressing PV inhibitory interneurons (625 nm) to isolate long-range CC inputs (455 nm). (B) Top Left: Photocurrent responses of a ChrimsonR-expressing PV cell to large-field red light stimulation when recorded in voltage-clamp (− 94 mV; 1000 ms pulse duration; tested with 9 LED intensities). Top Right: Summary graph shows light intensity and photocurrent response relationship measured for the peak and steady-state (SS) currents. Bottom Left: Loose-patch voltage recordings of a ChrimsonR-expressing PV cell while stimulating with 625 nm light (1000 ms pulse; tested with 9 LED intensities). Bottom Right: Summary graph shows light intensity and spike probability relationship measured for ChrimsonR-expressing PV cells. (C) Top Left: Voltage responses of a non-expressing L2/3 RS cell to large-field red light stimulation when recorded in current-clamp (−74 mV; 1000 ms pulse duration; tested with 9 LED intensities). Top Right: Summary graph shows light intensity and voltage response relationship measured for the peak hyperpolarization. Bottom Left: Voltage responses of a non-expressing L2/3 RS cell to intracellular positive current steps with (red) and without (black) PV cell photostimulation. Bottom Right: Summary graph showing spike rate with and without PV cell photostimulation (0.04 mW; 100 ms bins). (D) Left: Representative vS1-vM1 excitatory synaptic currents evoked by a pair of optical stimuli at 20 Hz (455nm, blue arrow, 0.5 ms) delivered with and without PV cell photostimulation (1000 ms; average of 10 trials each). Summary graph showing the paired-pulse ratio of the optically evoked CC responses for each cell with and without photostimulation of PV cells (0.04 mW). The red triangle represents the mean. Optogenetically silencing local excitatory cells did not result in short-term depression. (E) Left: Representative vS1-vM1 excitatory synaptic currents evoked by a 20 Hz optical train (455nm, blue arrow, 0.5 ms) delivered with and without PV cell photostimulation (1000 ms; average of 10 trials each). Summary graph showing EPSC amplitudes plotted as a function of stimulus number within 20 Hz trains. Optogenetically silencing local excitatory cells did not result in short-term depression. (F) In a subset of experiments, 50 μM APV was added to the bathing solution to reduce the responses of local excitatory cells by pharmacologically blocking non-linearities in the synaptic conductances associated with NMDA receptors. Current-clamp recording of a representative L2/3 RS cell during repetitive CC activation (20 Hz). Top: Control activity while holding the cell at a depolarized membrane potential (−74 mV). Note cells did not spike from their resting membrane potential with the LED intensities used for testing (Table S1). Bottom: the same cell and conditions except in the presence of 50 μM APV (overlay of 10 sweeps for each condition). The addition of APV suppressed excitatory cell excitability. (G) Left: Representative vM1-vS1 excitatory synaptic currents evoked by a pair of optical stimuli at 20 Hz delivered with and without APV (average of 10 trials each). Summary graph showing the paired-pulse ratio of the optically evoked responses for each cell with and without APV. The red triangle represents the mean. (H) Left: Representative vM1-vS1 excitatory synaptic currents evoked by a 20 Hz optical train delivered with and without APV (average of 10 trials each). Summary graph showing EPSC amplitudes plotted as a function of stimulus number within 20 Hz trains. Pharmacologically reducing local excitatory cell excitability did not result in short-term depression.

**Figure S5.**
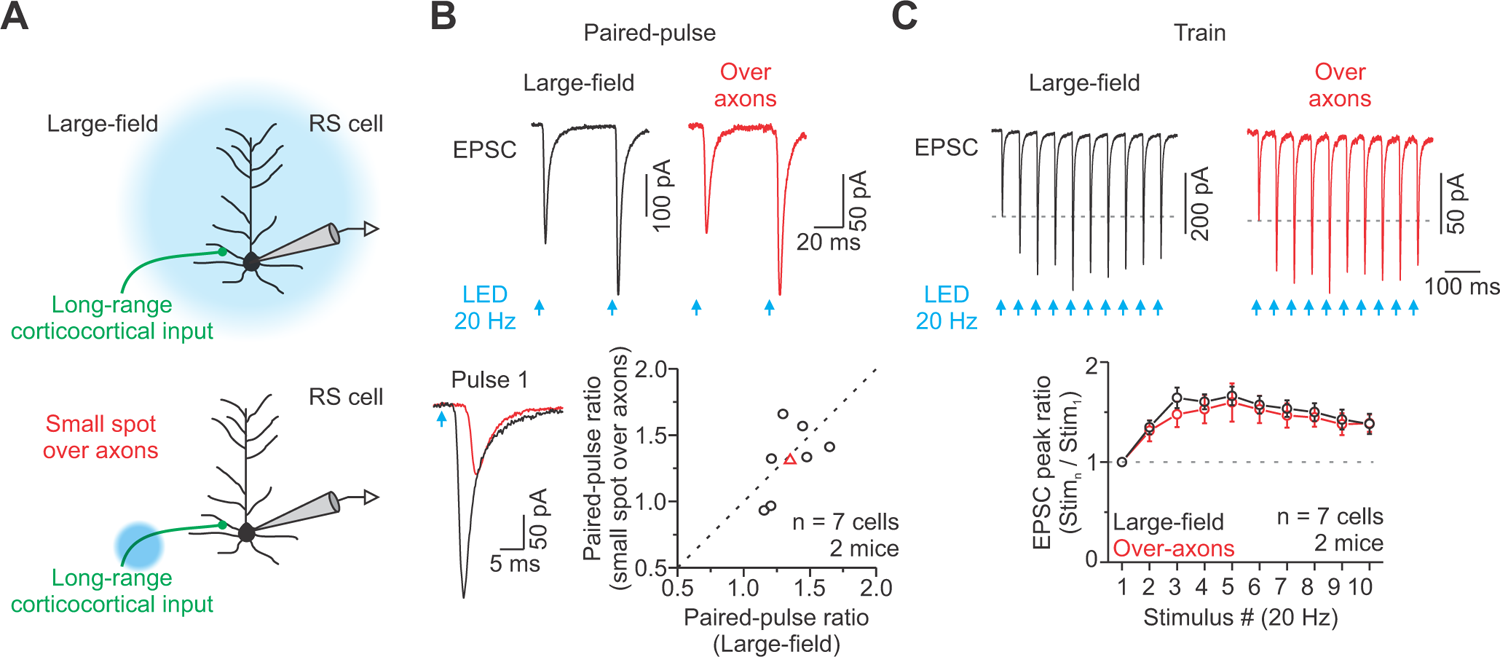
Large-field optically evoked short-term facilitation is similar to over-axon stimulation and not confounded by intracortical inputs. Related to Figure 1. (A) Recording schematic to test the performance of large-field optical stimulation. We recorded light-evoked excitatory responses recorded from an excitatory L2/3 RS cell during large-field illumination (standard approach) and over-axon stimulation (20 Hz, 0.5 ms pulse duration). Optical stimulation over-axons was achieved by restricting the light spot size and directing the light over the axons near white matter (800-1000 μm away). (B) Top: vM1 excitatory synaptic currents evoked by a pair of optical stimuli at 20 Hz (blue arrow, 0.5 ms) delivered using our standard approach (large-field) or over-axons (average of 14 trials each). Bottom Left: Axonally induced EPSCs showed a longer synaptic delay, confirming that spikes were initiated far from the recording pipette (Large-field: 2.1 ± 0.1 ms; Over-axon: 4.7 ± 0.3 ms). Bottom Right: Population data showing no difference in the paired-pulse ratio (PPR) for large-field versus over-axon optical stimulation (n = 7 cells from 2 mice; p = 0.71, Paired t-test). The red triangle represents the mean. (C) Excitatory synaptic currents evoked in the same neuron (shown in B, left) by a 20 Hz optical train delivered by our standard approach (large-field) or over-axons (average of 14 trials each). Short-term plasticity was similar for large-field and axon stimulation with 20 Hz trains (p = 0.24, Two-way ANOVA, stim. 2-10).

**Figure S6.**
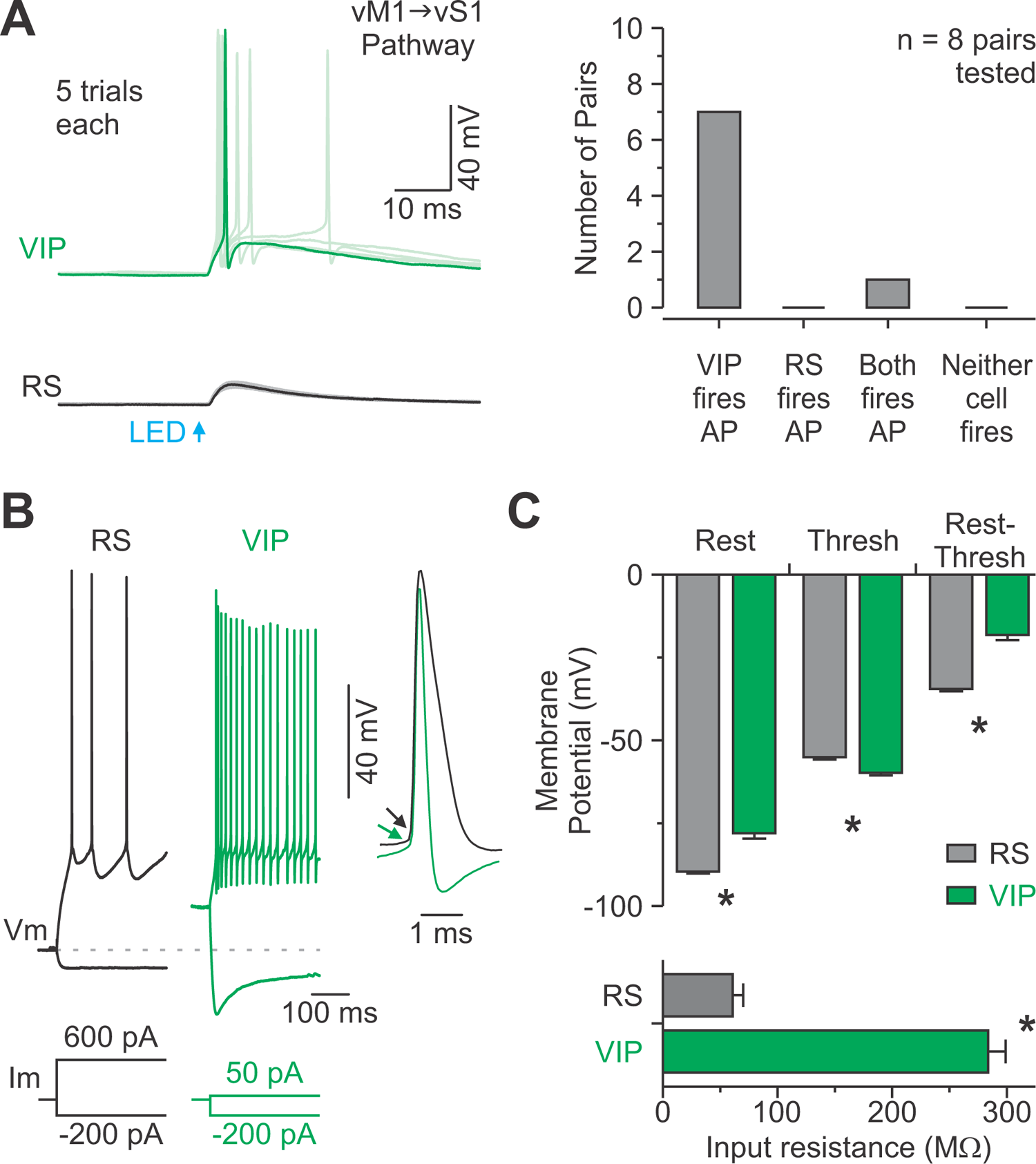
vM1-vS1 CC voltage responses were strongest in vS1 VIP cells despite similar synaptic current strengths in paired VIP-RS cell recordings due to their greater intrinsic excitability. Related to Figure 4. (A) Left: Paired VIP-RS cell recordings of CC responses in current-clamp at rest (5 trials shown). Action potentials can be seen ∼3.5 ms after CC photostimulation. Right: Summary of 8 cell pairs exhibiting spike responses to CC input (all current-clamp recordings at rest; max stimulus 32 mW). In all pairs tested, the VIP cell fired an action potential. (B) Responses of VIP-RS cell pair to injected current from rest (same cells as A). VIP cells were more depolarized at rest than RS cells. Voltage response to negative current step was larger in VIP than RS cells, reflecting lower membrane input resistance. A larger positive current was required to reach spike threshold in the RS than VIP cell (+450 pA versus +50 pA). Voltage thresholds for spiking were slightly lower in the VIP than RS cells. (C) Top: mean resting potential (Rest: p < 0.0001, Two-sample t-test), spike threshold (Thresh: p < 0.001, Two-sample t-test), and the differences between them (Rest-Thresh: p < 0.0001, Two-sample t-test; n = 10 pairs). Bottom: mean membrane input resistance (p < 0.001, Mann-Whitney U test; n = 10 pairs).

**Figure S7.**
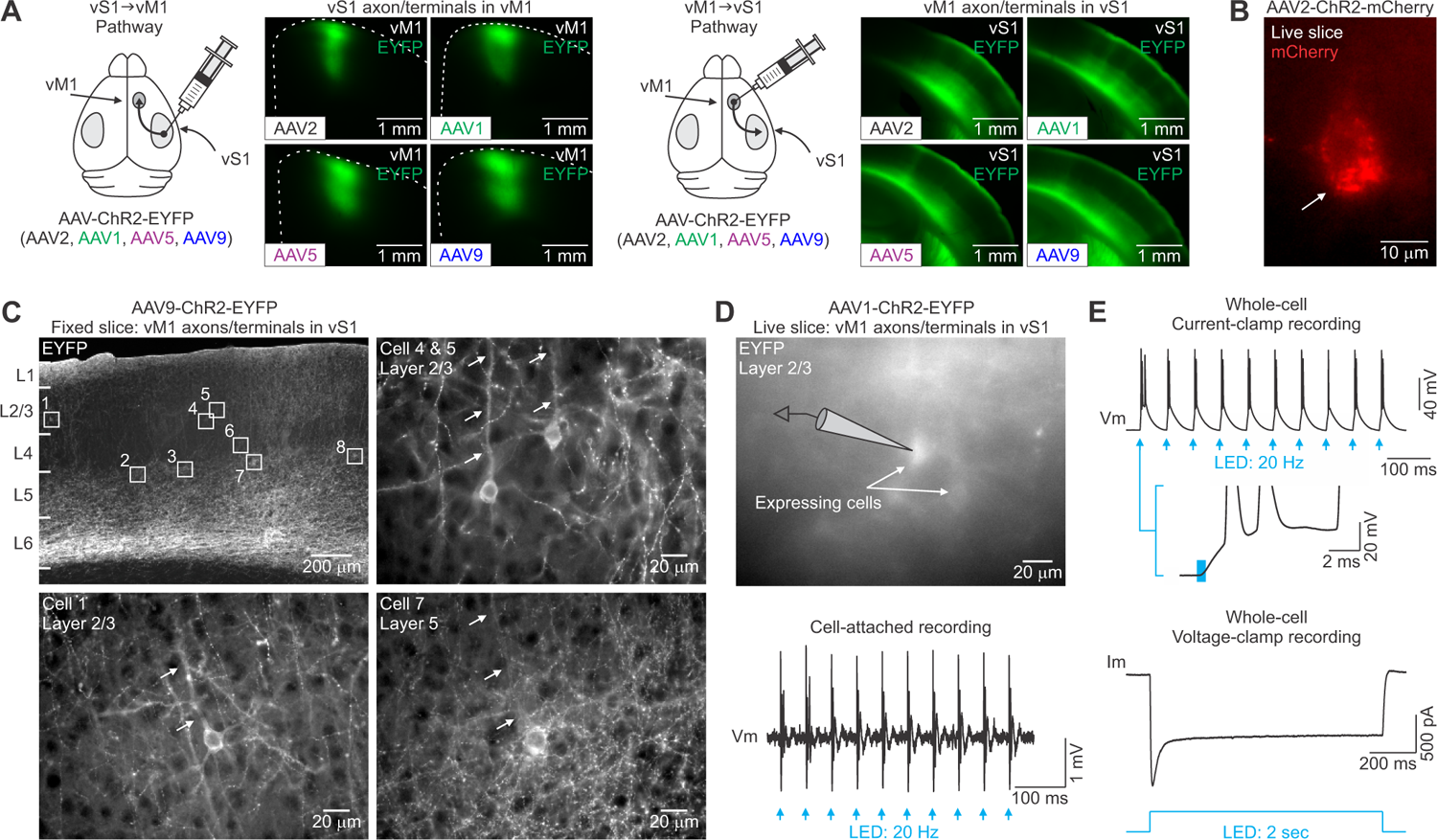
Retrograde viral transduction of CC circuits with certain AAVs and aggregation of ChR2-mCherry. Related to Figure 5. (A) Schematic showing virus injected into the vS1 (left) or vM1 (right) of mice *in vivo* (AAV2, AAV1, AAV5, or AAV9). We used the same vector and titer for each viral serotype (AAV-hsyn-ChR2(H124R)-EYFP). Live slice (300 μm) images showing vS1 (left) and vM1 (right) axon/terminal arbors expressing ChR2/EYFP after injection of each AAV. Expression levels were similar for all serotypes in both pathways (vS1-vM1: p = 0.81; AAV2, n = 6 mice; AAV1, n= 2 mice, AAV5, n = 3 mice; AAV9, n = 2 mice; vM1-vS1: p = 0.82; AAV2, n = 7 mice; AAV1, n= 5 mice, AAV5, n = 3 mice; AAV9, n = 7 mice, One-way ANOVA). (B) A high-magnification fluorescence image of a live 300 μm thick slice showing a representative ChR2-mCherry expressing neuron in vS1 (23 days after viral injection in vS1). Notice that ChR2-mCherry appears to be aggregated within the membrane or cytoplasm of the cell (white arrow). Similar results were observed in 3 mice. (C) Top left: Low-magnification fluorescence image of a 30-μm-thick section showing AAV9-driven expression of ChR2-EYFP in vM1 axons terminating in vS1 (20 days after virus injection in vM1), as well as retrograde viral expression in cortical neurons (boxes). High-magnification images from the same section show cortical cells in vS1 with EYFP-labeled somata and apical dendrites (arrows). We routinely observed signs of retrograde viral transduction of cortical neurons using AAV1, AAV5, and AAV9, but not AAV2 (n = 3 mice each). Most expressing cells had a pyramidal shape, characteristic RS physiology, and concentrated in L2/3 and L5a, where many CC cells populate (Harris and Shepherd 2015). (D) Top: Epifluorescence image of EYFP-expressing neurons in L2/3 of vS1 (live 300-μm-thick slice), 20 days after virus injection in vM1. Bottom: From the same cell shown above, voltage recordings of light-evoked responses in cell-attached mode (0.5 ms pulse duration, 10 pulses, 20 Hz, 32 mW or 18.0 mW/mm^2^). Spikes were evoked on each flash. (E) During whole-cell recording, responses of the same cell (shown in B) to a 20 Hz optical train of 0.5 ms light flashes (32 m or 18.0 mW/mm^2^W; blue arrows). Again, spikes were evoked on each flash. Membrane depolarization began almost immediately at flash onset (<0.5 ms). Bottom: Current recording of the same ChR2-expressing L2/3 cell to a 2 s LED stimulus (1.7 mW or 1.0 mW/mm^2^). The inward current initially peaked, then decreased to a sustained level within 100 ms. We obtained similar results in all 7 cells recorded after AAV1, AAV5, or AAV9 viral injections. Consistent with the anatomical work, less than 2% of all cells recorded after AAV2 injections showed signs of retrograde viral transduction (n = 8/527 cells).

**Figure S8.**
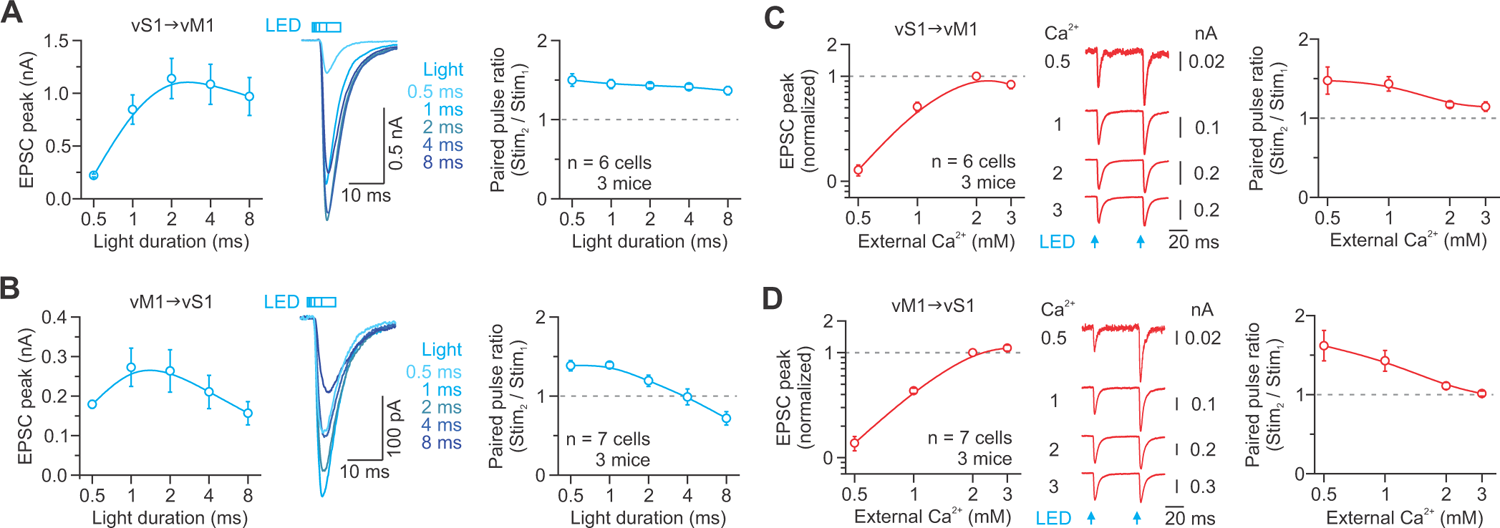
Effect of light duration and extracellular calcium on optically evoked short-term plasticity at CC synapses. Related to Figure 6. (A-B) Left: Population data showing the average EPSC amplitude of vS1-vM1 (A) and vM1-vS1 (B) synapses on L2/3 RS cells recorded in response to increasing LED pulse duration (different shades of blue: 0.5 – 8.0 ms). Middle: Representative EPSCs evoked with different LED pulse durations (average of 10-12 trials). Right: Population plot showing the peak paired-pulse ratio recorded in response to different LED pulse durations (50 ms interstimulus interval) (vS1-vM1: n = 6 cells from 3 mice; vM1-vS1: n = 7 cells from 3 mice). Increasing the LED pulse duration from 0.5 to 4 and 8 ms resulted in a significant decrease in paired-pulse facilitation for the vM1-vS1 pathway (p < 0.01, One-way ANOVA, with Bonferroni’s post-hoc test). Duration did not influence the peak paired-pulse ratio for the vS1-vM1 pathway (p = 0.53, One-way ANOVA). (C-D) Left: Population data showing the average normalized EPSC amplitude for vS1-vM1 (C) and vM1-vS1 (D) synapses on L2/3 RS cells recorded in different external Ca^2+^ concentrations. Data were normalized to the amplitude in 2 mM Ca^2+^. Middle: A pair of optically evoked EPSCs recorded in different external Ca^2+^ (50 ms interstimulus interval; average of 9-19 trials each). Right: Population plot showing the peak paired-pulse ratio recorded in different external Ca^2+^ (50 ms interstimulus interval) (vS1-vM1: n = 6 cells from 3 mice; vM1-vS1: n = 7 cells from 3 mice). Raising Ca^2+^ increased EPSC size and decreased paired-pulse facilitation.

**Figure S9.**
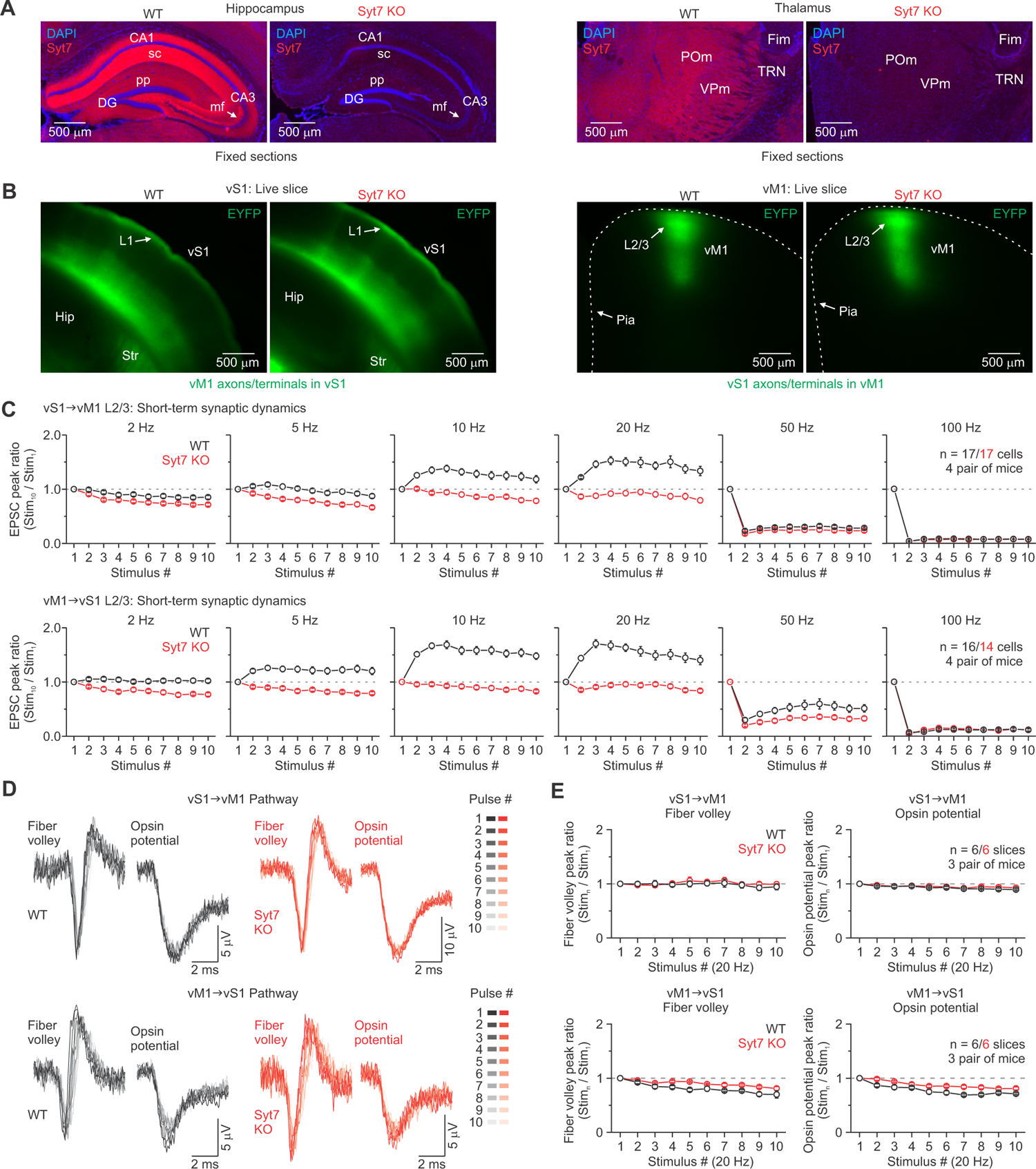
Facilitation is absent in Syt7 KO mice at multiple frequencies. Related to Figure 7. (A) Fluorescence images of the hippocampus (left) and thalamus (right) in a P116 WT and a Syt7 KO littermate mouse immunostained for Syt7 and counterstained with DAPI. In WT mice, Syt7 labeling is present in hippocampal and thalamic regions containing synapses known to facilitate, such as the Schaffer collateral synapses (sc), the mossy fiber synapses (mf), perforant path synapses (pp), thalamic reticular nucleus (TRN), ventral posterior medial nucleus (VPm), and posterior medial nucleus (POm). Thalamic regions are known to receive dense input from L6 corticothalamic cells, which facilitate. All regions lacked Syt7 expression in the KO animal. (B) Left: Epifluorescence images of live coronal slices (300 μm) showing ChR2/EYFP-expressing vM1 terminal arbors in vS1 of a P65 WT mouse and a P64 Syt7 KO littermate (23 and 22 days of expression). Right: Epifluorescence images of live coronal slices (300 μm) showing ChR2/EYFP-expressing vS1 terminal arbors in vM1 of a P44 WT mouse and a P43 Syt7 KO littermate (22 and 21 days of expression). Notice that the overall pattern of EYFP labeling in regions containing long-range vM1 or vS1 terminal arbors is similar in both WT and KO mice (n = 5 WT/KO pairs for vM1 synapses; n = 4 WT/KO pairs for vS1 synapses). This anatomical data suggests that the absence of Syt7 does not alter the structural organization of CC projections. Population plots of the average normalized peak synaptic responses evoked by 2-100 Hz optical trains (10 stimuli; 0.5 ms pulse duration) at the vS1-vM1 (top) and vM1-vS1 (bottom) synapses in slices prepared from WT and Syt7 KO littermates. In Syt7 KO mice, synaptic facilitation during trains was absent for both synapses at all frequencies tested. (D) ChR2 can reliably drive action potentials (APs) in CC axons from both WT and Syt7 KO animals. Overlay of the fiber volleys and opsin potentials recorded from the activation of the vS1-vM1 (top) and vM1-vS1 pathway (bottom) for both WT (black) and Syt7 KO mice (red). Responses were evoked by a 20 Hz train of optical stimuli (10 stimuli, 0.5 ms pulse duration, an average of ∼200 trials each). See Figure S2 for a description of how presynaptic APs (i.e., fiber volley) and opsin potentials were isolated from the recorded LFP. Light intensities were adjusted for each slice to the power needed to evoke a 200 pA EPSC in a neighboring L2/3 RS cell (average WT/KO LED powers: 1.4/1.1 mW or 0.8/0.6 mW/mm^2^ for vS1 axons in vM1, 12.7/6.1 mW or 7.2/3.2 mW/mm^2^ for vM1 axons in vS1). (E) Population data showing the normalized fiber volley amplitude and opsin potential amplitude during 20 Hz trains (n = 6/6 slices from 3 pairs of WT/KO mice vS1-vM1 pathway; n = 6/6 slices from 3 pairs of WT/KO mice vM1-vS1 pathway). Repetitive optical stimulation of ChR2-expressing axons/terminals resulted in similar fiber volley decay in both WT and KO animals. Overall, the decays were minimal for both animals in each pathway. Thus, the loss of short-term facilitation observed in the Syt7 KO animal (Figure 7) is not due to unreliable action potential generation (i.e., the number of axons recruited) during high-frequency stimulation. Values are represented as mean ± SEM.

---

**Table S1.** vM1 and vS1 synaptic responses in current-clamp and LED power information

| Cortex | Layer | EPSP Amplitude |  | Full-field LED Power |  |  |
| --- | --- | --- | --- | --- | --- | --- |
|  |  | EPSP peak (mv) | n cells | Total Power (mW) | Power Density (mW*mm<sup>2</sup>) | n cells |
| vS1 | L2/3 | 3.0 ± 0.3 | 41 | 12.8 ± 1.3 | 7.2 ± 0.7 | 61 |
| vS1 | L4 | 0.6 ± 0.2 | 2 | - | - | - |
| vS1 | L5 | 2.1 ± 0.4 | 17 | 19.3 ± 3.0 | 10.9 ± 1.7 | 22 |
| vS1 | L6 | 5.1 ± 0.3 | 35 | 4.8 ± 0.7 | 2.7 ± 0.4 | 18 |
| vM1 | L2/3 | 3.0 ± 0.2 | 31 | 1.7 ± 0.3 | 1.0 ± 0.2 | 54 |
| vM1 | L5 | 2.9 ± 0.4 | 19 | 2.9 ± 0.5 | 1.6 ± 0.4 | 24 |
| vM1 | L6 | 4.1 ± 0.6 | 15 | 10.8 ± 3.7 | 6.1 ± 2.1 | 13 |

**Table S2.** Layer boundaries based on layer specific Cre mouse lines. Related to **Figure 3**.

| Cortex | Layer | Cre line | Top boundaries |  | Bottom boundaries |  | *n*<br>slices/mice |
| --- | --- | --- | --- | --- | --- | --- | --- |
|  |  |  | Relative<br>depth | Absolute<br>depth (μm) | Relative<br>depth | Absolute<br>depth (μm) |  |
| vS1 | L4 | *Scnn1-Tg3-Cre* | 0.32 ± 0.01 | 409.3 ± 14.6 | 0.47 ± 0.01 | 608.5 ± 17.8 | 6/4 |
| vS1 | L5 | *Rbp4-Cre* | 0.48 ± 0.01 | 636.9 ± 15.2 | 0.72 ± 0.00 | 942.2 ± 16.5 | 13/7 |
| vS1 | L6 | *Ntsr1-Cre* | 0.69 ± 0.01 | 956.7 ± 12.2 | 0.97 ± 0.00 | 1343.2 ± 16.9 | 9/5 |
| vM1 | L4 | *Scnn1-Tg3-Cre* | - | - | - | - | 3/3 |
| vM1 | L5 | *Rbp4-Cre* | 0.30 ± 0.01 | 430.5 ± 15.2 | 0.54 ± 0.01 | 785.3 ± 16.5 | 10/5 |
| vM1 | L6 | *Ntsr1-Cre* | 0.70 ± 0.01 | 1053.6 ± 14.7 | 0.92 ± 0.01 | 1376.6 ± 17.3 | 8/4 |

- To characterize the pattern of Cre expression in each driver mouse we bred to the Ai14 tdTomato reporter line (Jackson 007908) - All measurements were based on live slice images - Relative depth is normalized to pia (0) and white matter (1) - No Cre expression was observed in vM1 of the *Scnn1-Tg3-Cre* mouse line

**Table S3.** Electrophysiological properties of wild type (WT) and Syt7 knockout (KO) neurons in layer 2/3 vS1. Related to **Figure 7**.

|  | WT |  | Syt7 KO |  | *P* | Test |
| --- | --- | --- | --- | --- | --- | --- |
|  | L2/3 RS cells |  | L2/3 RS cells |  |  |  |
|  | Mean ± s.e.m. | *n* cells / *n* mice | Mean ± s.e.m. | *n* cells / *n* mice |  |  |
| RMP (mV) | -89.7 ± 0.8 | 14 / 3 | -89.0 ± 0.9 | 15 / 3 | 0.643 | Mann-Whitney *U*-test |
| R<sub>m</sub> (MΩ) | 48.1 ± 2.6 | 14 / 3 | 46.4 ± 2.4 | 15 / 3 | 0.371 | Mann-Whitney *U*-test |
| τ<sub>m</sub>(ms) | 12.7 ± 0.5 | 14 / 3 | 12.1 ± 0.5 | 15 / 3 | 0.369 | two-sample *t*-test |
| C<sub>m</sub> (pF) | 273.7 ± 17.0 | 14 / 3 | 263.9 ± 8.1 | 15 / 3 | 0.599 | two-sample *t*-test |
| Rheobase (pA) | 292.5 ± 16.0 | 12 / 3 | 298.3 ± 23.3 | 12 / 3 | 0.838 | two-sample *t*-test |
| Spike threshold (mV) | -57.1 ± 0.4 | 12 / 3 | -57.1 ± 1.1 | 12 / 3 | 0.993 | two-sample *t*-test |
| Spike amplitude (mV) | 91.8 ± 0.9 | 12 / 3 | 92.6 ± 1.1 | 12 / 3 | 0.470 | Mann-Whitney *U*-test |
| Spike half-width (ms) | 0.72 ± 0.01 | 12 / 3 | 0.67 ± 0.01 | 12 / 3 | 0.012* | two-sample *t*-test |
| AHP amplitude (mV) | -14.5 ± 0.6 | 12 / 3 | -13.7 ± 1.0 | 12 / 3 | 0.458 | two-sample *t*-test |
| Max rate of rise (mV*ms) | 586.5 ± 15.6 | 12 / 3 | 621.8 ± 20.3 | 12 / 3 | 0.181 | two-sample *t*-test |
| Max rate of decay (mV*ms) | -99.9 ± 2.5 | 12 / 3 | -106.4 ± 2.7 | 12 / 3 | 0.091 | two-sample *t*-test |

• See methods for an explanation of how the electrophysiological parameters were defined/measured.
• All membrane potentials were corrected for a -14 mV liquid junction potential
• All statistical tests were two-tailed

## REFERENCES

1. Abbott LF, Regehr WG. 2004. Synaptic computation. Nature. 431:796–803.

2. Abbott LF, Varela JA, Sen K, Nelson SB. 1997. Synaptic depression and cortical gain control. Science. 275:220–224.

3. Adesnik H. 2018. Layer-specific excitation/inhibition balances during neuronal synchronization in the visual cortex. J Physiol. 596:1639–1657.

4. Agmon A, Connors BW. 1991. Thalamocortical responses of mouse somatosensory (barrel) cortex in vitro. Neuroscience. 41:365–379.

5. Andersen P, Sundberg SH, Sveen O, Wigstrom H. 1977. Specific long-lasting potentiation of synaptic transmission in hippocampal slices. Nature. 266:736–737.

6. Asrican B, Augustine GJ, Berglund K, Chen S, Chow N, Deisseroth K, Feng G, Gloss B, Hira R, Hoffmann C, Kasai H, Katarya M, Kim J, Kudolo J, Lee LM, Lo SQ, Mancuso J, Matsuzaki M, Nakajima R, Qiu L, Tan G, Tang Y, Ting JT, Tsuda S, Wen L, Zhang X, Zhao S. 2013. Next-generation transgenic mice for optogenetic analysis of neural circuits. Frontiers in neural circuits. 7:160.

7. Bastos AM, Usrey WM, Adams RA, Mangun GR, Fries P, Friston KJ. 2012. Canonical microcircuits for predictive coding. Neuron. 76:695–711.

8. Bastos AM, Vezoli J, Bosman CA, Schoffelen JM, Oostenveld R, Dowdall JR, De Weerd P, Kennedy H, Fries P. 2014. Visual Areas Exert Feedforward and Feedback Influences through Distinct Frequency Channels. Neuron.

9. Beierlein M, Gibson JR, Connors BW. 2003. Two dynamically distinct inhibitory networks in layer 4 of the neocortex. J Neurophysiol. 90:2987–3000.

10. Brown SP, Hestrin S. 2009. Intracortical circuits of pyramidal neurons reflect their long-range axonal targets. Nature. 457:1133–1136.

11. Bruno RM, Sakmann B. 2006. Cortex is driven by weak but synchronously active thalamocortical synapses. Science. 312:1622–1627.

12. Carvell GE, Miller SA, Simons DJ. 1996. The relationship of vibrissal motor cortex unit activity to whisking in the awake rat. Somatosens Mot Res. 13:115–127.

13. Castro-Alamancos MA. 2013. The motor cortex: a network tuned to 7-14 Hz. Frontiers in neural circuits. 7:21.

14. Cauller LJ, Clancy B, Connors BW. 1998. Backward cortical projections to primary somatosensory cortex in rats extend long horizontal axons in layer I. J Comp Neurol. 390:297–310.

15. Chakrabarti S, Kobayashi KS, Flavell RA, Marks CB, Miyake K, Liston DR, Fowler KT, Gorelick FS, Andrews NW. 2003. Impaired membrane resealing and autoimmune myositis in synaptotagmin VII-deficient mice. J Cell Biol. 162:543–549.

16. Chen JL, Carta S, Soldado-Magraner J, Schneider BL, Helmchen F. 2013. Behaviour-dependent recruitment of long-range projection neurons in somatosensory cortex. Nature. 499:336–340.

17. Cohen-Kashi Malina K, Mohar B, Rappaport AN, Lampl I. 2016. Local and thalamic origins of correlated ongoing and sensory-evoked cortical activities. Nature communications. 7:12740.

18. Collins DP, Anastasiades PG, Marlin JJ, Carter AG. 2018. Reciprocal Circuits Linking the Prefrontal Cortex with Dorsal and Ventral Thalamic Nuclei. Neuron. 98:366–379 e364.

19. Cossell L, Iacaruso MF, Muir DR, Houlton R, Sader EN, Ko H, Hofer SB, Mrsic-Flogel TD. 2015. Functional organization of excitatory synaptic strength in primary visual cortex. Nature. 518:399–403.

20. Covic EN, Sherman SM. 2011. Synaptic properties of connections between the primary and secondary auditory cortices in mice. Cerebral cortex. 21:2425–2441.

21. Crandall SR, Cruikshank SJ, Connors BW. 2015. A corticothalamic switch: controlling the thalamus with dynamic synapses. Neuron. 86:768–782.

22. Crandall SR, Govindaiah G, Cox CL. 2010. Low-threshold Ca2+ current amplifies distal dendritic signaling in thalamic reticular neurons. J Neurosci. 30:15419–15429.

23. Crandall SR, Patrick SL, Cruikshank SJ, Connors BW. 2017. Infrabarrels Are Layer 6 Circuit Modules in the Barrel Cortex that Link Long-Range Inputs and Outputs. Cell reports. 21:3065–3078.

24. Cruikshank SJ, Urabe H, Nurmikko AV, Connors BW. 2010. Pathway-specific feedforward circuits between thalamus and neocortex revealed by selective optical stimulation of axons. Neuron. 65:230–245.

25. D’Souza RD, Meier AM, Bista P, Wang Q, Burkhalter A. 2016. Recruitment of inhibition and excitation across mouse visual cortex depends on the hierarchy of interconnecting areas. eLife. 5.

26. Diamond ME, von Heimendahl M, Knutsen PM, Kleinfeld D, Ahissar E. 2008. ’Where’ and ‘what’ in the whisker sensorimotor system. Nat Rev Neurosci. 9:601–612.

27. Dong H, Shao Z, Nerbonne JM, Burkhalter A. 2004. Differential depression of inhibitory synaptic responses in feedforward and feedback circuits between different areas of mouse visual cortex. J Comp Neurol. 475:361–373.

28. Douglas RJ, Koch C, Mahowald M, Martin KA, Suarez HH. 1995. Recurrent excitation in neocortical circuits. Science. 269:981–985.

29. Engel AK, Fries P, Singer W. 2001. Dynamic predictions: oscillations and synchrony in top-down processing. Nat Rev Neurosci. 2:704–716.

30. Feldmeyer D, Lubke J, Sakmann B. 2006. Efficacy and connectivity of intracolumnar pairs of layer 2/3 pyramidal cells in the barrel cortex of juvenile rats. J Physiol. 575:583–602.

31. Feldmeyer D, Lubke J, Silver RA, Sakmann B. 2002. Synaptic connections between layer 4 spiny neurone-layer 2/3 pyramidal cell pairs in juvenile rat barrel cortex: physiology and anatomy of interlaminar signalling within a cortical column. J Physiol. 538:803–822.

32. Ferezou I, Haiss F, Gentet LJ, Aronoff R, Weber B, Petersen CC. 2007. Spatiotemporal dynamics of cortical sensorimotor integration in behaving mice. Neuron. 56:907–923.

33. Frandolig JE, Matney CJ, Lee K, Kim J, Chevee M, Kim SJ, Bickert AA, Brown SP. 2019. The Synaptic Organization of Layer 6 Circuits Reveals Inhibition as a Major Output of a Neocortical Sublamina. Cell reports. 28:3131–3143 e3135.

34. Friedman WA, Zeigler HP, Keller A. 2012. Vibrissae motor cortex unit activity during whisking. J Neurophysiol. 107:551–563.

35. Gabernet L, Jadhav SP, Feldman DE, Carandini M, Scanziani M. 2005. Somatosensory integration controlled by dynamic thalamocortical feed-forward inhibition. Neuron. 48:315–327.

36. Gil Z, Connors BW, Amitai Y. 1999. Efficacy of thalamocortical and intracortical synaptic connections: quanta, innervation, and reliability. Neuron. 23:385–397.

37. Gilbert CD, Li W. 2013. Top-down influences on visual processing. Nat Rev Neurosci. 14:350–363.

38. Gong S, Doughty M, Harbaugh CR, Cummins A, Hatten ME, Heintz N, Gerfen CR. 2007. Targeting Cre recombinase to specific neuron populations with bacterial artificial chromosome constructs. J Neurosci. 27:9817–9823.

39. Gunaydin LA, Yizhar O, Berndt A, Sohal VS, Deisseroth K, Hegemann P. 2010. Ultrafast optogenetic control. Nat Neurosci. 13:387–392.

40. Harris KD, Shepherd GM. 2015. The neocortical circuit: themes and variations. Nat Neurosci. 18:170–181.

41. Hass CA, Glickfeld LL. 2016. High-fidelity optical excitation of cortico-cortical projections at physiological frequencies. J Neurophysiol. 116:2056–2066.

42. Hempel CM, Hartman KH, Wang XJ, Turrigiano GG, Nelson SB. 2000. Multiple forms of short-term plasticity at excitatory synapses in rat medial prefrontal cortex. J Neurophysiol. 83:3031–3041.

43. Hill DN, Curtis JC, Moore JD, Kleinfeld D. 2011. Primary motor cortex reports efferent control of vibrissa motion on multiple timescales. Neuron. 72:344–356.

44. Hippenmeyer S, Vrieseling E, Sigrist M, Portmann T, Laengle C, Ladle DR, Arber S. 2005. A developmental switch in the response of DRG neurons to ETS transcription factor signaling. PLoS Biol. 3:e159.

45. Hoffer ZS, Hoover JE, Alloway KD. 2003. Sensorimotor corticocortical projections from rat barrel cortex have an anisotropic organization that facilitates integration of inputs from whiskers in the same row. J Comp Neurol. 466:525–544.

46. Hooks BM, Hires SA, Zhang YX, Huber D, Petreanu L, Svoboda K, Shepherd GM. 2011. Laminar analysis of excitatory local circuits in vibrissal motor and sensory cortical areas. PLoS Biol. 9:e1000572.

47. Hooks BM, Mao T, Gutnisky DA, Yamawaki N, Svoboda K, Shepherd GM. 2013. Organization of cortical and thalamic input to pyramidal neurons in mouse motor cortex. J Neurosci. 33:748–760.

48. Jackman SL, Beneduce BM, Drew IR, Regehr WG. 2014. Achieving high-frequency optical control of synaptic transmission. J Neurosci. 34:7704–7714.

49. Jackman SL, Regehr WG. 2017. The Mechanisms and Functions of Synaptic Facilitation. Neuron. 94:447–464.

50. Jackman SL, Turecek J, Belinsky JE, Regehr WG. 2016. The calcium sensor synaptotagmin 7 is required for synaptic facilitation. Nature. 529:88–91.

51. Jouhanneau JS, Kremkow J, Dorrn AL, Poulet JF. 2015. In Vivo Monosynaptic Excitatory Transmission between Layer 2 Cortical Pyramidal Neurons. Cell reports. 13:2098–2106.

52. Kaneko T, Caria MA, Asanuma H. 1994. Information-Processing within the Motor Cortex.2. Intracortical Connections between Neurons Receiving Somatosensory Cortical Input and Motor Output Neurons of the Cortex. J Comp Neurol. 345:172-184.

53. Kim J, Matney CJ, Blankenship A, Hestrin S, Brown SP. 2014. Layer 6 corticothalamic neurons activate a cortical output layer, layer 5a. J Neurosci. 34:9656–9664.

54. Kinnischtzke AK, Fanselow EE, Simons DJ. 2016. Target-specific M1 inputs to infragranular S1 pyramidal neurons. J Neurophysiol. 116:1261–1274.

55. Kinnischtzke AK, Simons DJ, Fanselow EE. 2014. Motor cortex broadly engages excitatory and inhibitory neurons in somatosensory barrel cortex. Cerebral cortex. 24:2237–2248.

56. Klapoetke NC, Murata Y, Kim SS, Pulver SR, Birdsey-Benson A, Cho YK, Morimoto TK, Chuong AS, Carpenter EJ, Tian Z, Wang J, Xie Y, Yan Z, Zhang Y, Chow BY, Surek B, Melkonian M, Jayaraman V, Constantine-Paton M, Wong GK, Boyden ES. 2014. Independent optical excitation of distinct neural populations. Nat Methods. 11:338–346.

57. Kleinfeld D, Ahissar E, Diamond ME. 2006. Active sensation: insights from the rodent vibrissa sensorimotor system. Curr Opin Neurobiol. 16:435–444.

58. Krupa DJ, Wiest MC, Shuler MG, Laubach M, Nicolelis MA. 2004. Layer-specific somatosensory cortical activation during active tactile discrimination. Science. 304:1989–1992.

59. Larkum M. 2013. A cellular mechanism for cortical associations: an organizing principle for the cerebral cortex. Trends Neurosci. 36:141–151.

60. Larkum ME, Nevian T, Sandler M, Polsky A, Schiller J. 2009. Synaptic integration in tuft dendrites of layer 5 pyramidal neurons: a new unifying principle. Science. 325:756–760.

61. Lee S, Carvell GE, Simons DJ. 2008. Motor modulation of afferent somatosensory circuits. Nat Neurosci. 11:1430–1438.

62. Lee S, Kruglikov I, Huang ZJ, Fishell G, Rudy B. 2013. A disinhibitory circuit mediates motor integration in the somatosensory cortex. Nat Neurosci. 16:1662–1670.

63. Lien AD, Scanziani M. 2013. Tuned thalamic excitation is amplified by visual cortical circuits. Nat Neurosci. 16:1315–1323.

64. Lin JY, Lin MZ, Steinbach P, Tsien RY. 2009. Characterization of engineered channelrhodopsin variants with improved properties and kinetics. Biophys J. 96:1803–1814.

65. Madisen L, Zwingman TA, Sunkin SM, Oh SW, Zariwala HA, Gu H, Ng LL, Palmiter RD, Hawrylycz MJ, Jones AR, Lein ES, Zeng H. 2010. A robust and high-throughput Cre reporting and characterization system for the whole mouse brain. Nat Neurosci. 13:133–140.

66. Manita S, Suzuki T, Homma C, Matsumoto T, Odagawa M, Yamada K, Ota K, Matsubara C, Inutsuka A, Sato M, Ohkura M, Yamanaka A, Yanagawa Y, Nakai J, Hayashi Y, Larkum ME, Murayama M. 2015. A Top-Down Cortical Circuit for Accurate Sensory Perception. Neuron. 86:1304–1316.

67. Mao T, Kusefoglu D, Hooks BM, Huber D, Petreanu L, Svoboda K. 2011. Long-range neuronal circuits underlying the interaction between sensory and motor cortex. Neuron. 72:111–123.

68. Markram H, Wang Y, Tsodyks M. 1998. Differential signaling via the same axon of neocortical pyramidal neurons. Proc Natl Acad Sci USA. 95:5323–5328.

69. McCormick DA, Connors BW, Lighthall JW, Prince DA. 1985. Comparative electrophysiology of pyramidal and sparsely spiny stellate neurons of the neocortex. J Neurophysiol. 54:782–806.

70. Naskar S, Qi J, Pereira F, Gerfen CR, Lee S. 2021. Cell-type-specific recruitment of GABAergic interneurons in the primary somatosensory cortex by long-range inputs. Cell reports. 34:108774.

71. O’Connor DH, Peron SP, Huber D, Svoboda K. 2010. Neural activity in barrel cortex underlying vibrissa-based object localization in mice. Neuron. 67:1048–1061.

72. Olsen SR, Bortone DS, Adesnik H, Scanziani M. 2012. Gain control by layer six in cortical circuits of vision. Nature. 483:47–52.

73. Petreanu L, Huber D, Sobczyk A, Svoboda K. 2007. Channelrhodopsin-2-assisted circuit mapping of long-range callosal projections. Nat Neurosci. 10:663–668.

74. Petreanu L, Mao T, Sternson SM, Svoboda K. 2009. The subcellular organization of neocortical excitatory connections. Nature. 457:1142–1145.

75. Petrof I, Viaene AN, Sherman SM. 2015. Properties of the primary somatosensory cortex projection to the primary motor cortex in the mouse. J Neurophysiol. 113:2400–2407.

76. Porter LL, White EL. 1983. Afferent and efferent pathways of the vibrissal region of primary motor cortex in the mouse. J Comp Neurol. 214:279–289.

77. Reyes A, Lujan R, Rozov A, Burnashev N, Somogyi P, Sakmann B. 1998. Target-cell-specific facilitation and depression in neocortical circuits. Nat Neurosci. 1:279–285.

78. Reyes A, Sakmann B. 1999. Developmental switch in the short-term modification of unitary EPSPs evoked in layer 2/3 and layer 5 pyramidal neurons of rat neocortex. J Neurosci. 19:3827–3835.

79. Rocco-Donovan M, Ramos RL, Giraldo S, Brumberg JC. 2011. Characteristics of synaptic connections between rodent primary somatosensory and motor cortices. Somatosens Mot Res. 28:63–72.

80. Sabatini BL, Regehr WG. 1999. Timing of synaptic transmission. Annu Rev Physiol. 61:521–542.

81. Sherman SM, Guillery RW. 2011. Distinct functions for direct and transthalamic corticocortical connections. J Neurophysiol. 106:1068–1077.

82. Somjen GG. 2002. Ion regulation in the brain: implications for pathophysiology. Neuroscientist. 8:254–267.

83. Sreenivasan V, Esmaeili V, Kiritani T, Galan K, Crochet S, Petersen CC. 2016. Movement Initiation Signals in Mouse Whisker Motor Cortex. Neuron. 92:1368–1382.

84. Stratford KJ, Tarczy-Hornoch K, Martin KA, Bannister NJ, Jack JJ. 1996. Excitatory synaptic inputs to spiny stellate cells in cat visual cortex. Nature. 382:258–261.

85. Takahashi N, Ebner C, Sigl-Glockner J, Moberg S, Nierwetberg S, Larkum ME. 2020. Active dendritic currents gate descending cortical outputs in perception. Nat Neurosci. 23:1277–1285.

86. Taniguchi H, He M, Wu P, Kim S, Paik R, Sugino K, Kvitsiani D, Fu Y, Lu J, Lin Y, Miyoshi G, Shima Y, Fishell G, Nelson SB, Huang ZJ. 2011. A resource of Cre driver lines for genetic targeting of GABAergic neurons in cerebral cortex. Neuron. 71:995–1013.

87. Tervo DG, Hwang BY, Viswanathan S, Gaj T, Lavzin M, Ritola KD, Lindo S, Michael S, Kuleshova E, Ojala D, Huang CC, Gerfen CR, Schiller J, Dudman JT, Hantman AW, Looger LL, Schaffer DV, Karpova AY. 2016. A Designer AAV Variant Permits Efficient Retrograde Access to Projection Neurons. Neuron. 92:372–382.

88. Tsodyks MV, Markram H. 1997. The neural code between neocortical pyramidal neurons depends on neurotransmitter release probability. Proc Natl Acad Sci USA. 94:719–723.

89. Turecek J, Jackman SL, Regehr WG. 2017. Synaptotagmin 7 confers frequency invariance onto specialized depressing synapses. Nature. 551:503–506.

90. Veinante P, Deschenes M. 2003. Single-cell study of motor cortex projections to the barrel field in rats. J Comp Neurol. 464:98–103.

91. Vijayan S, Hale GJ, Moore CI, Brown EN, Wilson M. 2010. Activity in the barrel cortex during active behavior and sleep. J Neurophysiol. 103:2074–2084.

92. Weiler N, Wood L, Yu J, Solla SA, Shepherd GM. 2008. Top-down laminar organization of the excitatory network in motor cortex. Nat Neurosci. 11:360–366.

93. White EL, DeAmicis RA. 1977. Afferent and efferent projections of the region in mouse SmL cortex which contains the posteromedial barrel subfield. J Comp Neurol. 175:455–482.

94. Williams SR, Mitchell SJ. 2008. Direct measurement of somatic voltage clamp errors in central neurons. Nat Neurosci. 11:790–798.

95. Xu NL, Harnett MT, Williams SR, Huber D, O’Connor DH, Svoboda K, Magee JC. 2012. Nonlinear dendritic integration of sensory and motor input during an active sensing task. Nature. 492:247–251.

96. Yamashita T, Pala A, Pedrido L, Kremer Y, Welker E, Petersen CC. 2013. Membrane potential dynamics of neocortical projection neurons driving target-specific signals. Neuron. 80:1477–1490.

97. Yang W, Carrasquillo Y, Hooks BM, Nerbonne JM, Burkhalter A. 2013. Distinct balance of excitation and inhibition in an interareal feedforward and feedback circuit of mouse visual cortex. J Neurosci. 33:17373–17384.

98. Young H, Belbut B, Baeta M, Petreanu L. 2021. Laminar-specific cortico-cortical loops in mouse visual cortex. eLife. 10.

99. Zagha E, Casale AE, Sachdev RN, McGinley MJ, McCormick DA. 2013. Motor cortex feedback influences sensory processing by modulating network state. Neuron. 79:567–578.

100. Zhang S, Xu M, Kamigaki T, Hoang Do JP, Chang WC, Jenvay S, Miyamichi K, Luo L, Dan Y. 2014. Selective attention. Long-range and local circuits for top-down modulation of visual cortex processing. Science. 345:660–665.

101. Zhang YP, Oertner TG. 2007. Optical induction of synaptic plasticity using a light-sensitive channel. Nat Methods. 4:139–141.

102. Zolnik TA, Ledderose J, Toumazou M, Trimbuch T, Oram T, Rosenmund C, Eickholt BJ, Sachdev RNS, Larkum ME. 2020. Layer 6b Is Driven by Intracortical Long-Range Projection Neurons. Cell reports. 30:3492–3505 e3495.

103. Zucker RS. 1999. Calcium- and activity-dependent synaptic plasticity. Curr Opin Neurobiol. 9:305–313.

104. Zucker RS, Regehr WG. 2002. Short-term synaptic plasticity. Annu Rev Physiol. 64:355–405.

## SUPPLEMENTAL REFERENCES

105. Crandall SR, Patrick SL, Cruikshank SJ, Connors BW. 2017. Infrabarrels Are Layer 6 Circuit Modules in the Barrel Cortex that Link Long-Range Inputs and Outputs. Cell reports. 21:3065–3078.

106. Cruikshank SJ, Urabe H, Nurmikko AV, Connors BW. 2010. Pathway-specific feedforward circuits between thalamus and neocortex revealed by selective optical stimulation of axons. Neuron. 65:230–245.

107. Harris KD, Shepherd GM. 2015. The neocortical circuit: themes and variations. Nat Neurosci. 18:170–181.

108. Hass CA, Glickfeld LL. 2016. High-fidelity optical excitation of cortico-cortical projections at physiological frequencies. J Neurophysiol. 116:2056–2066.

109. Lin JY, Lin MZ, Steinbach P, Tsien RY. 2009. Characterization of engineered channelrhodopsin variants with improved properties and kinetics. Biophys J. 96:1803–1814.

110. Petreanu L, Mao T, Sternson SM, Svoboda K. 2009. The subcellular organization of neocortical excitatory connections. Nature. 457:1142–1145.

